# Homeostatic active zone remodeling consolidates memories in the *Drosophila* mushroom body

**DOI:** 10.1101/2021.12.01.470602

**Authors:** Oriane Turrel, Niraja Ramesh, Marc J.F. Escher, Stephan J. Sigrist

## Abstract

Establishing a detailed understanding of how the distinct forms of synaptic plasticity spatio-temporally engage into the initial storage and subsequent consolidation of memories remains a fundamental challenge of neuroscience. In addition to the better understood postsynaptic plasticity, different forms of presynaptic plasticity are widely expressed in mammalian brains and apparently operate along Hebbian or homeostatic rules. Their behavioral relevance remains enigmatic, however. Lately, acute upregulation of active zone (AZ) scaffold protein BRP and release factor Unc13A via specific axonal transport factors were shown to mediate stable expression of presynaptic homeostatic plasticity (PHP) at *Drosophila* neuromuscular junctions (NMJs).

We here demonstrate that AZ scaling processes are specifically needed for stable expression of both, NMJ PHP as well as aversive olfactory mid-term memory within intrinsic neurons of the *Drosophila* mushroom body (MB). We first demonstrate that AZ upscaling via BRP is specifically needed for expression but not induction of NMJ homeostatic plasticity, thus establishing a direct temporal plasticity sequence of molecularly distinct AZ remodeling steps. Notably, when we reduced BRP and associated transport factors in MB intrinsic neurons, short-term memory persisted but robust deficits in stable memory expression for a few hours after conditioning were observed. In contrast, AZ release site protein RIM-BP affecting PHP induction was additionally needed for successful formation of short-term memory.

Taken together, our data establish a specific role of homeostatic presynaptic long-term plasticity for memory consolidation. Such homeostatic refinement processes might well be needed to successfully integrate and display synaptic engrams constituting intermediary term memories.

## INTRODUCTION

Synapses are key sites of information processing and of storage in the brain. The synaptic transmission strength is not hardwired but adapts during synaptic plasticity to provide adequate input-output relationships and maintain or restore transmission when compromised and to store information^1–7^. However, there is still a fundamental gap in our understanding of how dynamic changes of synapse performance intersect with circuit operation and, consequently, define behavioral states. This, to a great extent, is due to the inherent complexity of synaptic plasticity mechanisms, operating across many timescales (sub-second to days) and using a rich spectrum of both pre- and postsynaptic molecular and cellular mechanisms. In the context of learning and memory in rodents, Hebbian modifications are meant to alter and introduce positive feedback into networks to mark specific subsets of synapses for memory trace formation^8, 9^. In contrast, homeostatic plasticity mechanisms, which operate via negative feedbacks, are believed to compensate for such changes driving away from the equilibrium of synaptic strength and thus to constrain neuronal activity levels^10^. Homeostatic plasticity processes are observed from invertebrates through humans^11^, but are mechanistically well accessible at the neuromuscular junction (NMJ) of *Drosophila* larvae^11^. Here, fast presynaptic homeostatic plasticity (PHP) can be triggered by the application of glutamate receptor blocker Philanthotoxin-433 (PhTx; specifically blocking GluRIIA sub-unit containing receptors), and precisely counterbalances this postsynaptic glutamate receptor disturbance to enhance presynaptic neurotransmitter release. NMJ PHP increases both the release probability for docked SVs at existing release sites and also the number of functional release sites^12–14^. The presynaptic active zones (AZs) where synaptic vesicles (SVs) get released are covered by complex protein scaffolds, composed of a conserved set of extended proteins. Thereby, discrete “nanoscopic” release sites for SVs, defined by the ELKS/BRP localizing the critical unc13 family protein Unc13A in defined spatial and biochemical relations to the release triggering AZ Ca^2+^ channels, might be added to existing active zones^15, 16^.

Concerning a system where to test the role of such homeostatic plasticity processes, the Mushroom Body (MB) is an associative center in invertebrate brains meant to initially form and store memories. Presynaptic plasticity at the *Drosophila* MB intrinsic Kenyon cell (KCs) synapses is considered critical for the initialization of olfactory memory formation^17, 18^. Concerning the molecular synaptic plasticity processes driving MB memory consolidation less is known. Notably, the active zone architecture, including the exact nanoscale spacing between the BRP/Unc13 release machinery and the active zone central Ca^2+^ channels, is likely present across all *Drosophila* synapses including KC derived AZs ^19–21^.

We here test the principal relevance of presynaptic plasticity components concerning their role in MB olfactory memory formation and consolidation. Thus, we identify a mechanistic sequence at homeostatically remodeling NMJ AZs. Consistent with previous observations^22^, we find that in response to paired (but not unpaired) olfactory conditioning, BRP/Unc13A levels increased over the Mushroom Body formation for a few hours. Concerning NMJ PHP, AZ organizer protein BRP as well as the Arl8 GTPase (critical for trafficking of BRP-AZ-precursors) were essential for stable expression (measured at 30 min after PhTx application) but not induction (measured at 10 min). Post-developmental MB KCs-specific reduction of either BRP or Arl8 in turn let initial learning unaffected, but severely attenuated hours range consolidated mid-term memory. Unc104/IMAC and Aplip1, additional BRP-complex relevant transport components, resulted in very similar phenotypes. In contrast, removal of RIM-BP, essential for PHP induction, abrogated learning “already”. Thus, homeostatic plasticity processes involving structural AZ remodeling seem to play a role for the consolidation of learned information in MB KSs, illustrating how homeostatic plasticity can promote the successful post-learning implementation of learned information.

### Results

How does synaptic integration and plasticity steer network function in control of animal behavior? This question has been pressing in the neurosciences for decades. We here test the role of discrete presynaptic plasticity regulators and components in the induction and subsequent expression (“consolidation”) of aversive olfactory memories, thought to critically depend on presynaptic plasticity processes of MB KCs synapses.

### Active zone upscaling via BRP needed for expression but not induction of NMJ homeostatic plasticity

Previously, NMJ synapses were shown to increase BRP/Unc13A staining levels upon 10 minutes of PhTx treatment^15^ when quantified with confocal microscopy. Thus, a plasticity-related molecular remodeling of AZs seemingly is part of this form of homeostatic plasticity (see scheme in Fig. 1A; we hereafter refer to this BRP/Unc13A as “presynaptic scaling”). To mechanistically dissect PHP under longer time scales, previous analyses^23^ used a chromosomally encoded *glurIIA* receptor mutant which chronically lacked GluRIIA receptors. In order to more directly address whether indeed a temporal sequence of functional plasticity steps might underly PHP, we for this study extended our analysis from the standard post 10 minutes (min) to post 30 min PhTx application (Fig. 1B,C). In control *w1118* flies, the reduction of postsynaptic sensitivity evidenced by mEPSP amplitude reduction was still present at 30 min (Fig. 1B,C, *w1118*), and PHP also persisted to 30 min, obvious in an about two-fold increase of quantal content (Fig. 1C, *w1118*). Indeed, quantal content increases tended to be even higher at 30 min than at 10 min (Fig. 1B,C, *w1118*). In contrast, *brp* mutants, still displaying PHP at 10 minutes^23, 24^ after PhTx treatment, were blocked in their ability to maintain PHP after 30 minutes of PhTx application (Fig. 1C, *Brp*). Thus, BRP plays a crucial role specifically in the stable expression but not induction of PHP at NMJ active zones.

**Figure 1.**
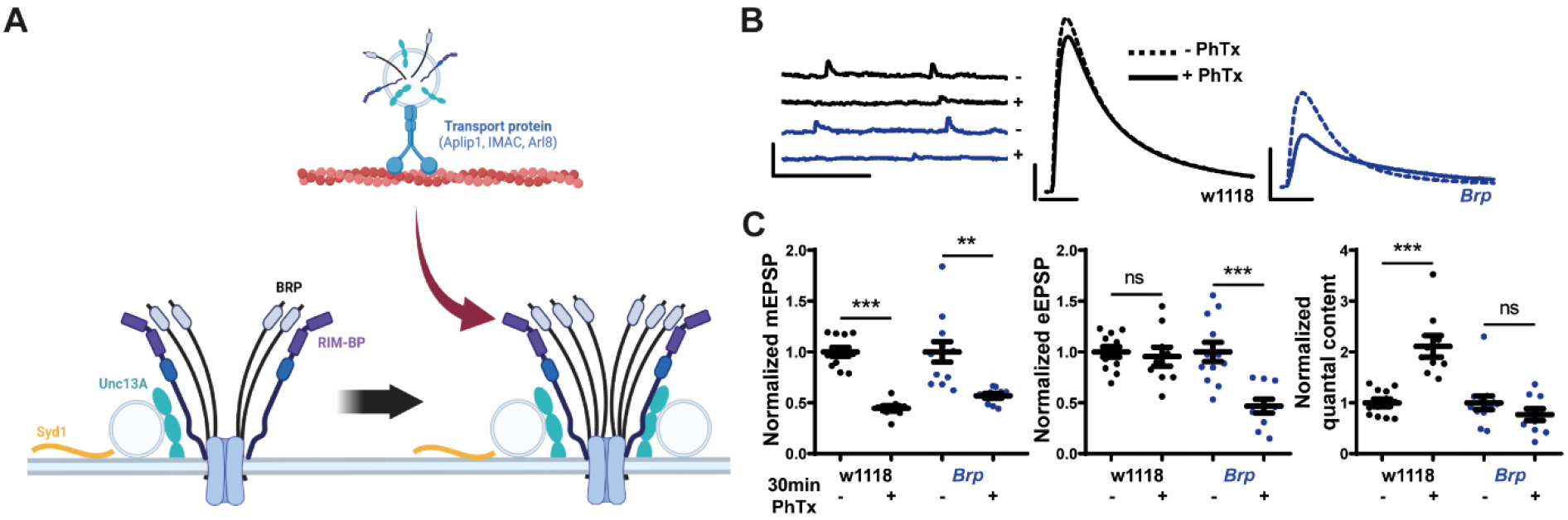
BRP is needed for long-term functional plasticity in larva. A. Schematic representation of the plasticity-related molecular remodeling of active zones at the NMJ. The main protein BRP form a ring around the Ca^2+^ Cac channel with RIM-BP. Synaptic vesicles are anchored to the membrane close to the BRP/RIM-BP/Cac channels by Unc13A and Syd1. Transport proteins, such as Aplip1, IMAC and Arl8, can bring new material to the AZ to reinforce it. B. Representative traces of mEPSP and eEPSP in wild-type *w1118* (black) and *brp*-null mutant (blue) before (- PhTx; continuous line) and after 30 min of PhTx (+ PhTx; dashed line) treatment. Scale bar: eEPSP 10 mV, 10 ms; mEPSP 5 mV, 500 ms. C. Quantifications of mEPSP amplitude, eEPSP amplitude and quantal content in PhTx-treated wild-type and *brp*-null mutant cells normalized on the same measurement obtained without PhTx for each genotype.

### Mushroom body active zones: BRP/Unc13A upscaling upon paired aversive olfactory conditioning

Recent work addressed mechanisms of presynaptic plasticity using the so-called STAR system^25^ to label MB KC-specific BRP expression triggered from the endogenous *brp* locus after induced recombination and introducing a FLAG-TAG^22^. When tested specifically in MB γ-neurons, this work uncovered a transient upregulation of BRP specifically upon paired olfactory conditioning. In order to also be able to analyze Unc13A, meant to be a major effector of presynaptic plasticity, we stained brains of adult wild-type (*w^1118^*) flies 1 h or 3 h after a single cycle of aversive conditioning for BRP (via Nc82 monoclonal), Syd1 and Unc13A, and quantified staining levels within the MB lobes (Fig. 2A). Indeed, animals displayed increases for BRP and even more pronounced for Unc13A at 1 h after training. While BRP levels plateaued 3 h after conditioning, Unc13A levels increased further towards 3 h (Fig. 2A). Syd1 levels, irresponsive to PhTx triggered homeostatic plasticity at NMJ synapses^23^, showed a significant increase when quantified in the MB lobes synapses only 3 h after aversive training (Fig. 2A).

**Figure 2.**
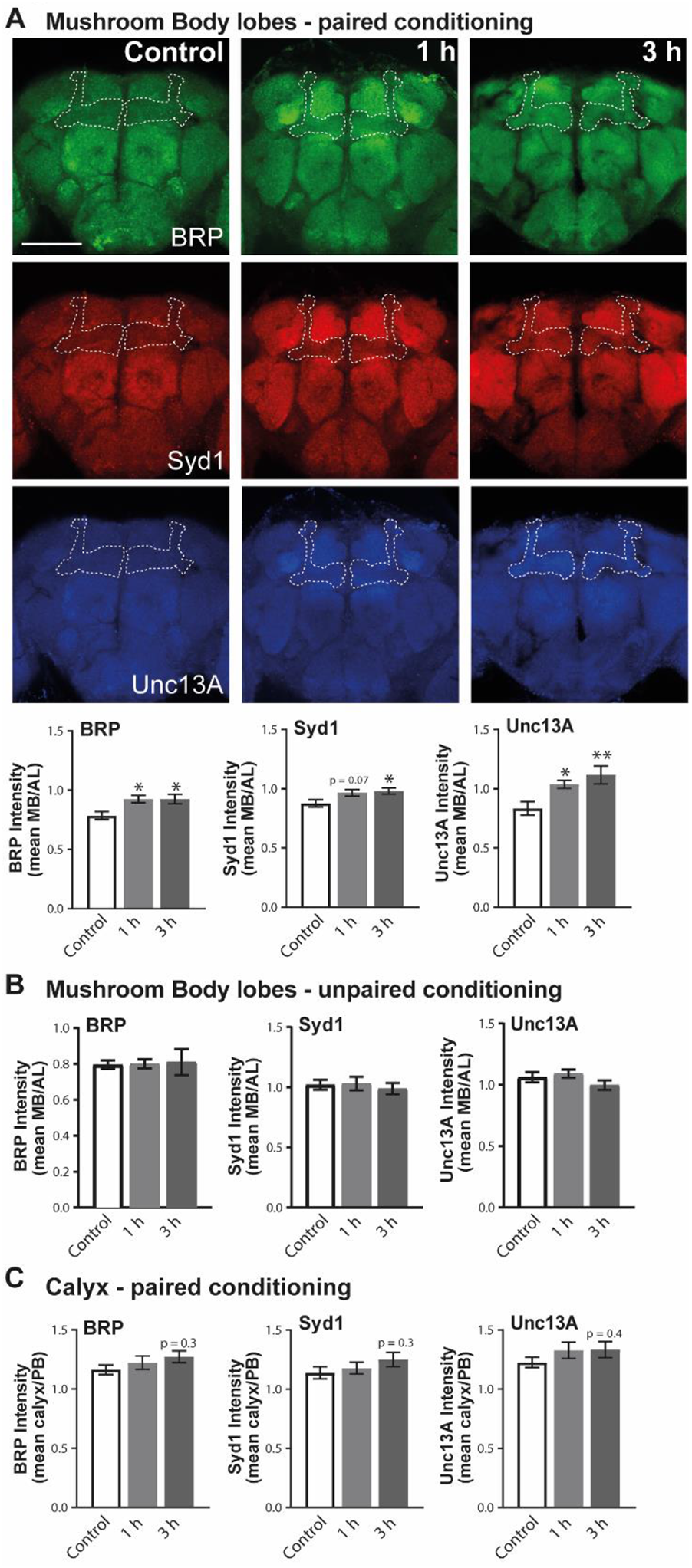
Memory formation leads to an increase of several AZ proteins indicating a reinforcement of the AZ structure. A. Representative confocal images for the staining. Quantification of BRP^nc82^, Syd1 and Unc13A staining intensity in MB lobes of *w1118* flies 1 h and 3 h after paired conditioning (BRP ^nc82^: *F*(_3,80_) = 5.215, *p* = 0.0075, *n* ≥ 26; *post hoc* Tukey’s multiple comparisons test, *Control* vs *1 h* \**p* < 0.05, *Control* vs *3 h* \**p* < 0.05; Syd1: *F*(_3,80_) = 3.857, *p* = 0.0253, *n* ≥ 26; *post hoc* Tukey’s multiple comparisons test, *Control* vs *1 h* p > 0.05, *Control* vs *3 h* \**p* < 0.05; Unc13A: *F*(_3,51_) = 6.258, *p* = 0.0038, *n* = 17; *post hoc* Tukey’s multiple comparisons test, *Control* vs *1 h* \**p* < 0.05, *Control* vs *3 h* \*\**p* < 0.01). Scale bar: 50 µm. B. Quantification of BRP ^nc82^, Syd1 and Unc13A staining intensity in MB lobes of *w1118* flies 1 h and 3 h after unpaired conditioning (BRP ^nc82^: *F*(_3,47_) = 0.1087, p = 0.8972, *n* ≥ 15; Syd1: *F*(_3,47_) = 0.6202, *p* = 0.5425, *n* ≥ 15; Unc13A: *F*(_3,47_) = 2.262, *p* = 0.1161, *n* ≥ 15). C. Quantification of BRP ^nc82^, Syd1 and Unc13A staining intensity in calyx of *w1118* flies 1 h and 3 h after paired conditioning (BRP ^nc82^: *F*(_3,72_) = 1.266, *p* = 0.2885, *n* ≥ 23; Syd1: *F*(_3,72_) = 1.136, *p* = 0.3270, *n* ≥ 23; Unc13A: *F*(_3,50_) = 1.004, *p* = 0.3740, *n* ≥ 16).

To verify whether these results were learning and memory relevant, meaning whether they were indeed specific to an association of the unconditioned stimulus (US, electric shock) and the particular odor (conditioned stimulus, CS), we also used an unpaired conditioning protocol. Here, shocks were presented before the odor, which should not lead to an aversive association of US and CS and consequently no learning or memory formation. Notably, unpaired conditioning did not change the intensities of BRP, Syd1 or Unc13A (Fig. 2B). Moreover, in the calyx of the MB no intensity changes were observed after paired conditioning (Fig. 2C), suggesting that the changes are at least particularly pronounced in the MB lobes and thus the presynaptic active zones of KCs might well be particularly enriched. It should be considered here, however, that MB lobe staining also contains contributions of other neuronal populations, particularly dopaminergic neurons.

### Post-development reduction of BRP in Kenyon cells attenuates mid- but not short-term memories

We next addressed whether homeostatic active zone plasticity mechanisms would be relevant for the formation of olfactory short-term memory formation (“learning”) and/or for longer lasting memories. As mentioned above, complete absence of BRP at larval NMJs still allows for the induction of homeostatic plasticity^23^ but abrogates stable expression of this plasticity form (^23, 24^, Fig. 1B,C). In contrast, active zone protein RIM-BP was needed for PHP at 10 minutes after PhTx onset^23^ “already”.

We here took care to not interfere with the principal organization of MB KC synapses, and at the same time to decrease protein levels to interfere with efficient plastic presynaptic scaling. Thus, we restricted the expression of the RNA interference (RNAi) constructs used to only the adult, post-developmental stage. Here, we took advantage of the TARGET system^26^, which entails a temperature sensitive Gal80 inhibitor (Gal80^ts^) to block the transcriptional activity of Gal4 at low temperature (18°C). At high temperature (29°C), however, the Gal80^ts^ protein is denatured and consequently the Gal4 activity inhibition lifted. In order to restrict the expression of the RNAi to the adult MB lobes, we combined *tub-Gal80^ts^* with the KCs generic *OK107-Gal4* line (*Gal80^ts^;OK107*).

First, we tested whether our post-developmental mRNA knockdown (KD) strategy would result in a scoreable reduction of the respective protein levels in the MB lobes. Indeed, after 5 days of temperature shift induction, flies expressing the RNAi-BRP^B3-C8^ ^27^ in the adult MB lobes showed an around 30% decrease of BRP intensity in the MB lobes, but no changes in the co-probed RIM-BP or Syd1. Reciprocally, flies expressing RNAi-RIM-BP in the adult MB lobes had an around 30% decrease of RIM-BP intensity in the MB lobes but no changes in BRP or Syd1 (Fig. 3A, see Supplementary Fig. 3A for representative images). Thus, we obviously could provoke a moderate but specific downregulation of both BRP and RIM-BP protein in the MB KCs.

**Figure 3.**
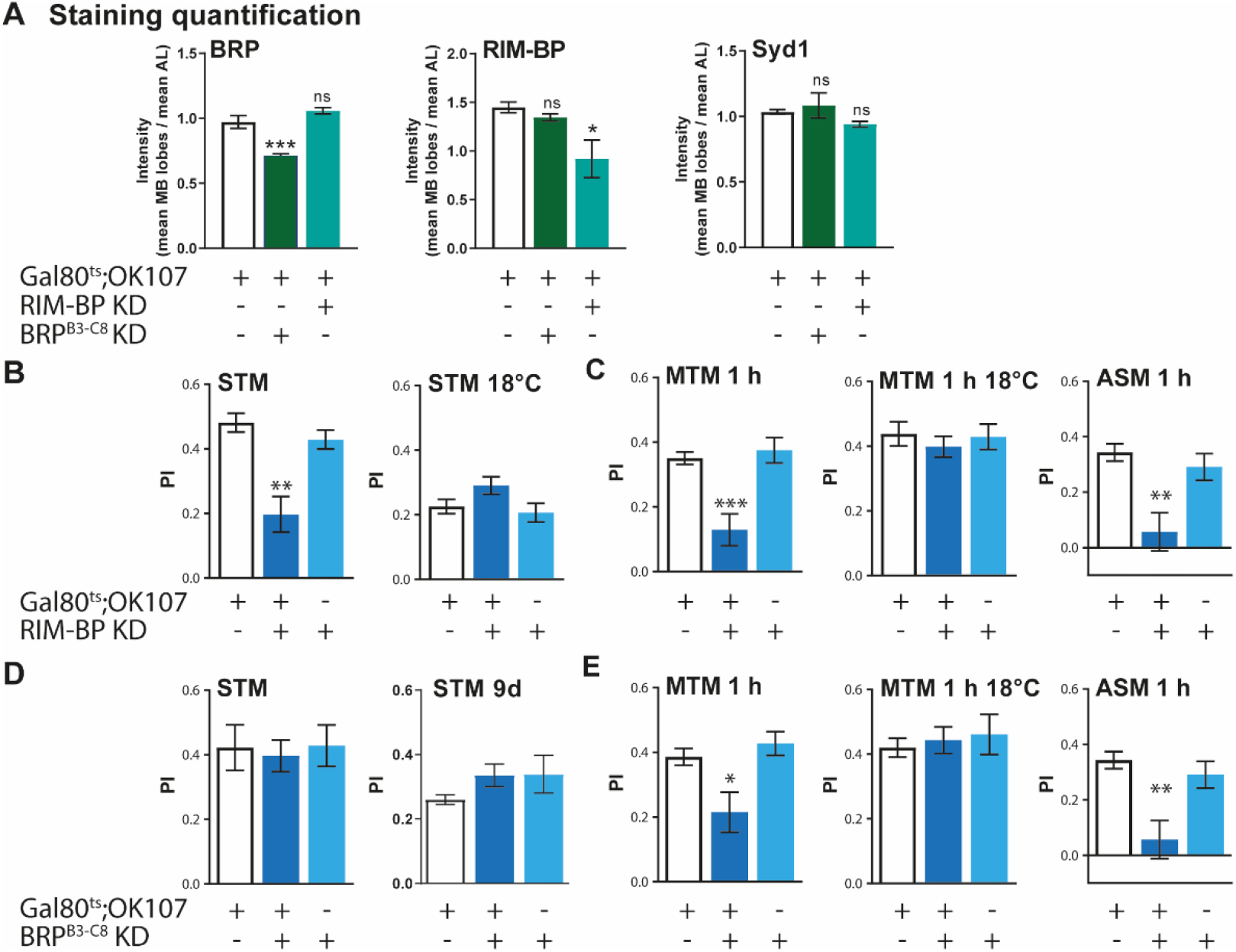
RIM-BP and BRP are needed in the adult MB lobes for MTM 1 h. A. Quantification of BRP ^nc82^, RIM-BP and Syd1 staining intensity in MB lobes of flies expressing RNAi-RIM-BP and RNAi-BRP^B3-C8^ in adult MB lobes (BRP ^nc82^: *F*(_3,13_) = 32.65, *p* < 0.0001, *n* ≥ 4; *post hoc* Tukey’s multiple comparisons test, *Gal80^ts^;OK107/+* vs *Gal80^ts^;OK107/RNAi-BRP^B3-C8^* \*\*\**p* < 0.001, *Gal80^ts^;OK107/+* vs *Gal80^ts^;OK107/RNAi-RIM-BP p* > 0.05; RIM-BP: *F*(_3,13_) = 4.504, *p* = 0.0403, *n* ≥ 4; *post hoc* Tukey’s multiple comparisons test, *Gal80^ts^;OK107/+* vs *Gal80^ts^;OK107/RNAi-RIM-BP* \**p* < 0.05, *Gal80^ts^;OK107/+* vs *Gal80^ts^;OK107/RNAi-BRP^B3-C8^ p* > 0.05; Syd1: *F*(_3,14_) = 1.312, *p* = 0.3084, *n* ≥ 4). B. Flies expressing RNAi-RIM-BP in the adult MB lobes exhibit lower STM compared to the genetic control (*F*(_3,24_) = 14.29, *p* = 0.0001, *n* = 8; *post hoc* Tukey’s multiple comparisons test, *Gal80^ts^;OK107/+* vs *Gal80^ts^;OK107/RNAi-RIM-BP* \*\*\**p* < 0.001, +/*RNAi-RIM-BP* vs *Gal80^ts^;OK107/RNAi-RIM-BP* **p < 0.01). Without induction, those flies have normal STM (*F*(_3,20_) = 2.926, *p* = 0.0809, *n* ≥ 6). C. *Gal80^ts^;OK107/RNAi-RIM-BP* flies present a deficit of MTM 1 h (*F*(_3,37_) = 12.63, *p* < 0.0001, *n* ≥ 12; *post hoc* Tukey’s multiple comparisons test, *Gal80^ts^;OK107/+* vs *Gal80^ts^;OK107/RNAi-RIM-BP* \*\*\**p* < 0.001, +/*RNAi-RIM-BP* vs *Gal80^ts^;OK107/RNAi-RIM-BP* ***p < 0.001). Without induction, those flies have normal MTM 1 h (*F*(_3,38_) = 0.3512, *p* = 0.7063, *n* ≥ 12). Flies expressing RNAi-RIM-BP in the adult MB lobes exhibit a deficit of ASM 1 h (*F*(_3,37_) = 8.581, *p* = 0.0010, *n* ≥ 12; *post hoc* Tukey’s multiple comparisons test, *Gal80^ts^;OK107/+* vs *Gal80^ts^;OK107/RNAi-RIM-BP* \*\**p* < 0.01, +/*RNAi-RIM-BP* vs *Gal80^ts^;OK107/RNAi-RIM-BP* **p < 0.01). D. Flies expressing RNAi-BRP^B3-C8^ have normal STM after 5 d of induction (*F*(_3,17_) = 0.07892, *p* = 0.9245, *n* ≥ 5) and also after 9 d of induction (*F*(_3,24_) = 1.217, *p* = 0.3162, *n* = 8). E. *Gal80^ts^;OK107/RNAi-BRP^B3-C8^* flies present a deficit of MTM 1 h (*F*(_3,60_) = 6.501, *p* = 0.0029, *n* = 20; *post hoc* Tukey’s multiple comparisons test, *Gal80^ts^;OK107/+* vs *Gal80^ts^;OK107/RNAi-BRP^B3-C8^* \**p* < 0.05, *+/RNAi-BRP^B3-C8^* vs *Gal80^ts^;OK107/RNAi-BRP^B3-C8^* \*\**p* < 0.01). Without induction, those flies have normal MTM 1 h (*F*(_3,29_) = 0.2266, *p* = 0.7988, *n* ≥ 9). Flies expressing RNAi-BRP^B3-C8^ in the adult MB lobes have a defect of ASM 1 h (*F*(_3,60_) = 7.915, *p* = 0.0009, *n* = 20; *post hoc* Tukey’s multiple comparisons test, *Gal80^ts^;OK107/+* vs *Gal80^ts^;OK107/RNAi-BRP^B3-C8^* \**p* < 0.05, *+/RNAi-BRP^B3-C8^* vs *Gal80^ts^;OK107/RNAi-BRP^B3-C8^* \*\*\**p* < 0.001).

We then tested the memory of these two KD genotypes using the classical associative aversive olfactory conditioning. Here, groups of flies are successively exposed to two distinct odors, the first one being associated to electric shocks. *Drosophila* presents different phases of associative aversive olfactory memory^17, 18, 28^: Short-Term Memory (STM) is measured immediately after a single conditioning round, whereas Mid-Term Memory (MTM) is typically evaluated in between 1 h and 3 h after training. Notably, when measuring MTM, two memory components, Anesthesia-Sensitive Memory (ASM), which is erased by anesthesia through cold shock, and Anesthesia-Resistant Memory (MT-ARM), can be discriminated. As a precondition for testing olfactory learning and memory, we tested whether our KDs would affect the ability of the flies to avoid electric shocks and olfactory acuity to the odors of flies. Importantly, our KD genotypes still displayed normal perception of the conditioning stimuli (Table 1), allowing their subsequent behavioral testing.

**Table 1.** Shock reactivity and olfactory acuity of flies expressing RNAi-RIM-BP or RNAi-BRP^B3-C8^ in the adult MB lobes. Data are shown as means ± SEM. After 5 d of induction, flies show normal shock reactivity (RNAi-RIM-BP: *F(_3,22_)* = 0.1296, *p* = 0.8792, *n* ≥ 7; RNAi-BRP^B3-C8^: *F(_3,24_)* = 2.263, *p* = 0.1288, *n* = 8) and normal olfactory acuity for octanol (RNAi-RIM-BP: *F(_3,36_)* = 1.391, *p* = 0.2630, *n* = 12; RNAi-BRP^B3-C8^: *F(_3,24_)* = 0.8075, *p* = 0.4593, *n* = 8) and methycyclohexanol (RNAi-RIM-BP: *F(_3,36_)* = 1.976, *p* = 0.1547, *n* = 12; RNAi-BRP^B3-C8^: *F(_3,24_)* = 1.373, *p* = 0.2752, *n* = 8).

| Genotype | Shock reactivity | Olfactory acuity |  |
| --- | --- | --- | --- |
|  |  | Octanol | Methylcyclohexanol |
| *Gal80<sup>ts</sup>;OK107/+* | 0.6243 ± 0.09976 | 0.4983 ± 0.05912 | 0.7075 ± 0.04990 |
| *Gal80<sup>ts</sup>;OK107/RNAi-RIM-BP* | 0.5957 ± 0.06841 | 0.6492 ± 0.07711 | 0.5392 ± 0.06307 |
| *+/RNAi-RIM-BP* | 0.5675 ± 0.06930 | 0.5758 ± 0.05316 | 0.6917 ± 0.08155 |
| *Gal80<sup>ts</sup>;OK107/+* | 0.8950 ± 0.02765 | 0.5088 ± 0.08943 | 0.5725 ± 0.1076 |
| *Gal80<sup>ts</sup>;OK107/RNAi-BRP<sup>B3-C8</sup>* | 0.7950 ± 0.04892 | 0.6488 ± 0.06768 | 0.6313 ± 0.07527 |
| *+/RNAi-BRP<sup>B3-C8</sup>* | 0.7763 ± 0.04739 | 0.5425 ± 0.08516 | 0.4188 ± 0.09523 |

We thus turned to testing aversive STM and MTM. After 5 days of induction at 29°C, flies expressing a RNAi directed against RIM-BP in the adult MB lobes showed a significant defect of STM (Fig. 3B). MTM measured at 1 h was significantly reduced as well, obviously due to decreased ASM (Fig. 3C) while ARM was not affected (Supplementary Fig. 3B; note that ARM scores are still rather low at 1 h after conditioning). Importantly, these deficits were not observed when flies were incubated at 18°C (where the Gal80^ts^ suppressor is stable), proving that the memory deficits observed were indeed caused via RNAi induction in the adult MB lobes (Fig. 3B for STM and Fig. 3C for MTM 1 h). In contrast to the RIM-BP interference, flies expressing a BRP directed RNAi in the adult MB lobes displayed normal STM (Fig. 3D). We wondered whether this reflected a still too little reduction of BRP levels. Thus, we also tested STM after 9 (instead of 5) days of induction (Fig. 3D), with the same result however. Notably, in contrast to STM, MTM scores measured 1 h after conditioning were significantly affected by BRP KD. This deficit was specifically due to decreased ASM (Fig. 3E), while ARM remained stable (Supplementary Fig. 3B). Again, 18°C degree controls did not show any memory deficits when compared to their cognate controls (Fig. 3E), proving that our effects are truly mediated by lifting the blockade of RNAi construct expression.

We so far used *OK107-Gal4* as a pan-MB line. We wondered whether similar results concerning effect size and quality of the MTM component would be retrieved when further restricting our manipulation to a relevant KC subset. Thus, we took advantage of the *c739-Gal4* driver line^29^, restricting our manipulations to the α/β MB lobes known to be critical for olfactory aversive mid-term memories^18^. Indeed, upon expression of RNAis directed against RIM-BP or BRP, MTM-specific deficits comparable in size to the effects provoked via pan-MB manipulation were observed (Supplementary Fig. 3C,D; also see Supplementary Table). Thus, our principal conclusion, that the BRP scaffold protein is crucial for stable expression but not the induction of NMJ homeostatic active zone plasticity and at the same time specifically needed for MTM but not STM, is seemingly robust against restricting to relevant subpopulations of MB intrinsic neurons called KCs.

### Active zone scaffold transport machinery specifically needed for mid-term memory consolidation

So far, our data suggest that a moderate post-developmental deprivation of the active zone remodeling factor BRP results in a specific deficit of mid-term “consolidated” memory. As said above, the overall reduction of BRP steady state levels as measured via confocal microscopy were only moderately reduced (Fig. 3A). RNA interference should deprive the BRP-encoding mRNA pool. As the BRP protein once formed is supposedly rather stable (Sigrist and Bhukel, unpublished), we suspect that the new synthesis of BRP and its subsequent axonal transport might be rate-limiting for AZ remodeling (and consequently MTM formation).

Thus, we went on testing whether indeed transport factors critical for the effective assembly and transport of BRP-containing AZ precursor material might also specifically link to mid-term memory formation. We connected this to a further analysis whether indeed the NMJ PHP phenotypes would again agree to the same logic, meaning whether a specific need for 30 min stable PHP expression would match with a specific requirement for 1 h MTM.

We hereby intended to test different transport proteins involved in specific subsequent parts of the transport of cargo to the AZs: Arl8 is involved in the first step during the assembly of the cargo^30^, then IMAC, a kinesin involved in the actual transport along microtubules filaments^31^, and finally, Aplip1 an adaptor protein between specific vesicles and kinesin^32^. In this regard, we started with the small GTPase Arl8, which takes a crucial role in the assembly of transportable active zone precursor complexes^31^. Absence of Arl8 leads to an accumulation of AZ precursors already in motor neuron cell bodies and provokes a reduction of BRP at NMJ terminals (Fig. 5A). Loss of Arl8 precludes the expression of PHP in a *glurIIA* mutant background^30^, as said a genetic constellation taken as a surrogate of long-term depression of pharmacological GluRIIA blockade via PhTx. We directly tested *arl8*-null larvae concerning induction and expression of NMJ PHP, as well as AZ remodeling via BRP staining. As expected, PhTx-triggered BRP increase was abolished in the *arl8* mutants (Fig. 5A) and at the same time, after 10 min of PhTx treatment, *arl8* mutants showed normal PHP, with mEPSPs amplitude being decreased but eEPSPs amplitude remaining stable due to a robust quantal content increase (Fig. 5B). After 30 min of PhTX, instead of a robust homeostatic increase of quantal content as in controls, no change compared to PhTx-untreated controls was observed (Fig. 5C). This clear distinction concerning a requirement for PHP induction (10 min) versus expression (30 min) is remarkable.

**Figure 4.**
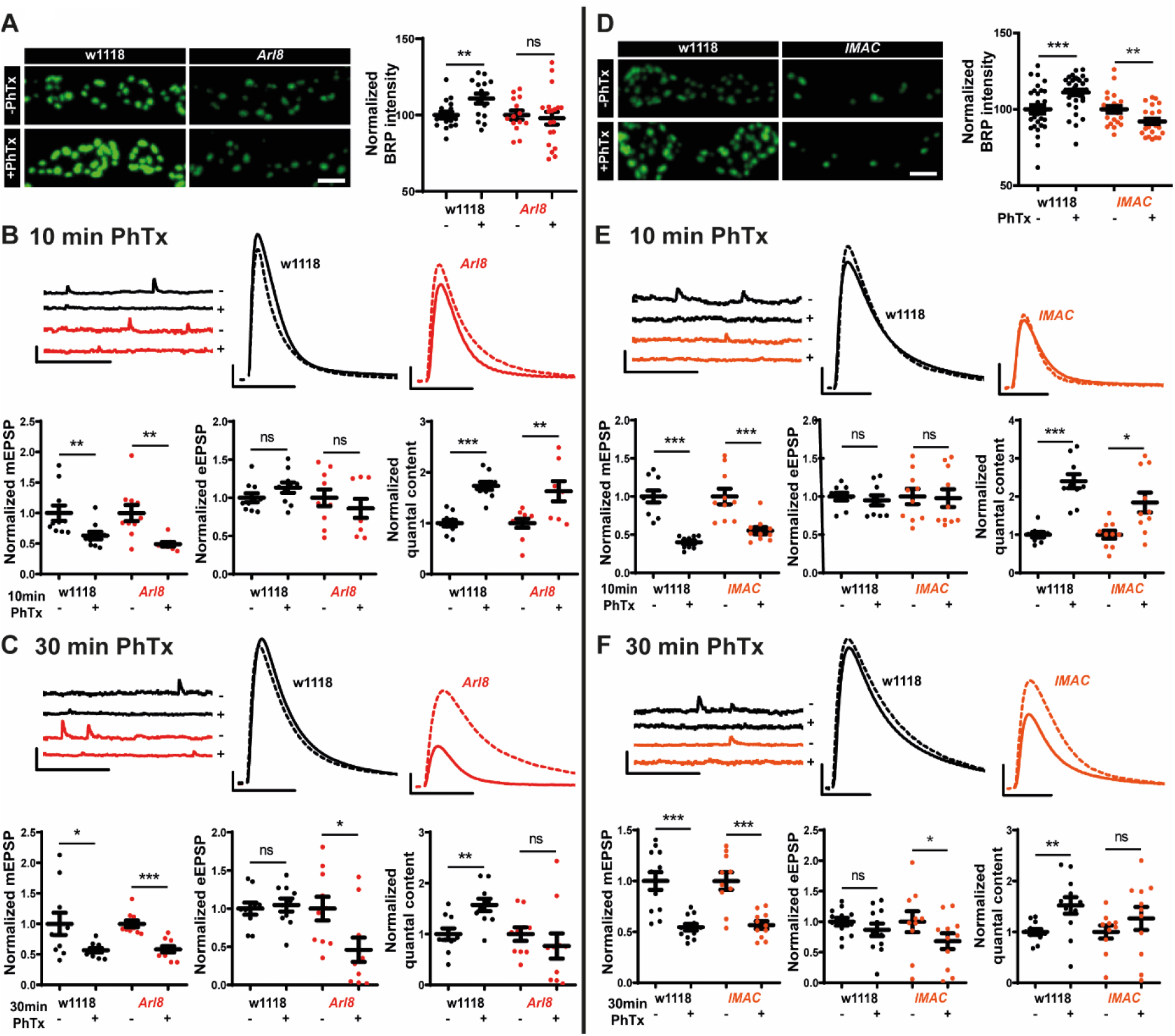
Transport proteins Arl8 and IMAC are both needed for 30 min functional plasticity in larva. A. Representative confocal images of muscle 4 NMJs of abdominal segment 2-5 from 3^rd^ instar larvae from wild-type *w1118* (left) and *arl8*-null mutant (right) NMJs labelled with BRP^nc82^ without (- PhTx) and after 10 min PhTx (+ PhTx) treatment and quantification of normalized BRP intensity. Scale bar: 2 µm. B. Representative traces of mEPSP and eEPSP in wild-type *w1118* (black) and *arl8*-null mutant (red) before (- PhTx; continuous line) and after 10 min of PhTx (+ PhTx; dashed line) treatment. Quantifications of mEPSP amplitude, eEPSP amplitude and quantal content in PhTx-treated wild-type and *arl8*-null mutant cells normalized on the same measurement obtained without PhTx for each genotype. Scale bar: eEPSP 10 mV, 10 ms; mEPSP 5 mV, 500 ms. C. Representative traces of mEPSP and eEPSP in wild-type *w1118* (black) and *arl8*-null mutant (red) before (- PhTx; continuous line) and after 30 min of PhTx (+ PhTx; dashed line) treatment. Quantifications of mEPSP amplitude, eEPSP amplitude and quantal content in PhTx-treated wild-type and *arl8*-null mutant cells normalized on the same measurement obtained without PhTx for each genotype. Scale bar: eEPSP 10 mV, 10 ms; mEPSP 5 mV, 500 ms. D. Representative confocal images of muscle 4 NMJs of abdominal segment 2-5 from 3^rd^ instar larvae from wild-type *w1118* (left) and *imac*-null mutant (right) NMJs labelled with BRP^nc82^ without (- PhTx) and after 10 min PhTx (+ PhTx) treatment and quantification of normalized BRP intensity. Scale bar: 2 µm. E. Representative traces of mEPSP and eEPSP in wild-type *w1118* (black) and *imac*-null mutant (orange) before (- PhTx; continuous line) and after 10 min of PhTx (+ PhTx; dashed line) treatment. Quantifications of mEPSP amplitude, eEPSP amplitude and quantal content in PhTx-treated wild-type and *imac*-null mutant cells normalized on the same measurement obtained without PhTx for each genotype. Scale bar: eEPSP 10 mV, 10 ms; mEPSP 5 mV, 500 ms. F. Representative traces of mEPSP and eEPSP in wild-type *w1118* (black) and *imac*-null mutant (orange) before (- PhTx; continuous line) and after 30 min of PhTx (+ PhTx; dashed line) treatment. Quantifications of mEPSP amplitude, eEPSP amplitude and quantal content in PhTx-treated wild-type and *imac*-null mutant cells normalized on the same measurement obtained without PhTx for each genotype. Scale bar: eEPSP 10 mV, 10 ms; mEPSP 5 mV, 500 ms.

**Figure 5.**
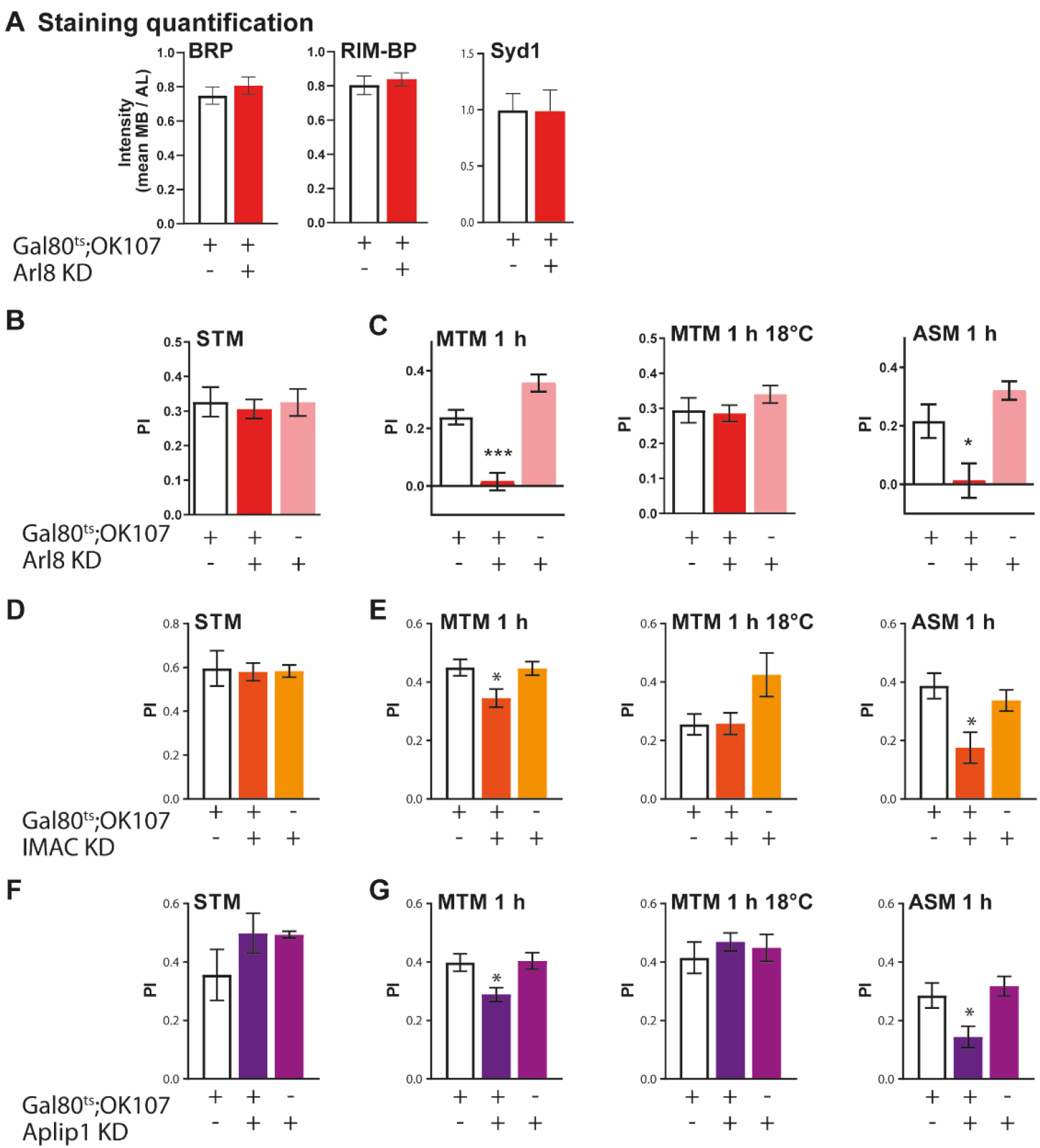
Transport proteins sustain specifically MTM in adult MB lobes. A. Quantification of BRP ^nc82^, RIM-BP and Syd1 staining intensity in MB lobes of flies expressing RNAi-Arl8 in adult MB lobes (BRP ^nc82^: *t-test*, *p* = 0.8200, *n* = 4; RIM-BP: *t-test*, *p* = 0.5811, *n* = 4; Syd1: *t-test*, *p* = 0.9546, *n* ≥ 9). B. After induction, *Gal80^ts^;OK107/RNAi-Arl8* have normal STM (*F*(_3,19_) = 0.1073, *p* = 0.8989, *n* ≥ 6). C. Flies expressing RNAi-Arl8 in the adult MB lobes present a deficit of MTM 1 h (*F*(_3,25_) = 34.17, *p* < 0.0001, *n* ≥ 7; *post hoc* Tukey’s multiple comparisons test, *Gal80^ts^;OK107/+* vs *Gal80^ts^;OK107/RNAi-Arl8* \*\*\**p* < 0.001, *+/RNAi-Arl8* vs *Gal80^ts^;OK107/RNAi-Arl8* \*\*\**p* < 0.001, *Gal80^ts^;OK107/+* vs *+/RNAi-Arl8* \**p* < 0.05), with a deficit of ASM 1 h (*F*(_3,24_) = 9.887, *p* = 0.0009, *n* ≥ 7; *post hoc* Tukey’s multiple comparisons test, *Gal80^ts^;OK107/+* vs *Gal80^ts^;OK107/RNAi-Arl8* \**p* < 0.05, *+/RNAi-Arl8* vs *Gal80^ts^;OK107/RNAi-Arl8* \*\*\**p* < 0.001). Without induction, those flies present normal MTM 1 h (*F*(_3,25_) = 1.195, *p* = 0.3215, *n* ≥ 7). D. Flies expressing RNAi-IMAC in adult MB lobes have normal STM (*F*(_3,20_) = 0.02769, *p* = 0.9727, *n* ≥ 5). E. *Gal80^ts^;OK107/RNAi-IMAC* flies present a MTM 1 h deficit (*F*(_3,51_) = 4.667, *p* = 0.0141, *n* = 17; *post hoc* Tukey’s multiple comparisons test, *Gal80^ts^;OK107/+* vs *Gal80^ts^;OK107/RNAi-IMAC* \**p* < 0.05, *+/RNAi-IMAC* vs *Gal80^ts^;OK107/RNAi-IMAC* \**p* < 0.05). Without induction, those flies have normal MTM 1 h (*F*(_3,24_) = 3.447, *p* = 0.0508, *n* = 8). Flies expressing RNAi-IMAC in adult MB lobes have a lower ASM 1 h compared to the genetic controls (*F*(_3,51_) = 6.095, *p* = 0.0044, *n* = 17; *post hoc* Tukey’s multiple comparisons test, *Gal80^ts^;OK107/+* vs *Gal80^ts^;OK107/RNAi-IMAC* \*\**p* < 0.01, *+/RNAi-IMAC* vs *Gal80^ts^;OK107/RNAi-IMAC* \**p* < 0.05). F. Flies expressing RNAi-Aplip1 in adult MB lobes have normal STM (*F*(_3,13_) = 1.236, *p* = 0.3315, *n* ≥ 3). G. Flies expressing RNAi-Aplip1 in adult MB lobes present a deficit of MTM 1 h (*F*(_3,52_) = 5.508, *p* = 0.0070, *n* ≥ 17; *post hoc* Tukey’s multiple comparisons test, *Gal80^ts^;OK107/+* vs *Gal80^ts^;OK107/RNAi-Aplip1* \**p* < 0.05, +/*RNAi-Aplip1* vs *Gal80^ts^;OK107/RNAi-Aplip1* *p < 0.05). Without induction, those flies have normal MTM 1 h (*F*(_3,24_) = 0.3712, *p* = 0.6943, *n* = 8). *Gal80^ts^;OK107/RNAi-Aplip1* flies exhibit a defect of ASM 1 h (*F*(_3,52_) = 5.983, *p* = 0.0047, *n* ≥ 17; *post hoc* Tukey’s multiple comparisons test, *Gal80^ts^;OK107/+* vs *Gal80^ts^;OK107/RNAi-Aplip1* \**p* < 0.05, +/*RNAi-Aplip1* vs *Gal80^ts^;OK107/RNAi-Aplip1* **p < 0.01).

Despite the suggestive character of these findings, it should be said that Arl8, apart from active zone precursor transport, executes other functions, e.g. the transport of lysosomes^30, 33, 34^. We thus sought to generate independent proof for our conclusion. In *C. elegans*, Arl8 acts in conjunction with the anterogradely kinesin Unc104/IMAC^31^. We thus analyzed *imac* mutants in the identical manner as *arl8* mutants (Fig. 4D-F), finding the exact same “phenotype profile”, though with a somewhat decreased effect size. Concretely, under PhTx, absence of BRP increase (PhTx 10 min, Fig. 4D), unperturbed PHP induction (PhTx 10 min, Fig. 4E), but defective PHP expression (PhTx 30 min, Fig. 4F).

In conclusion, Arl8 provides a more severe deficit whereas IMAC offers a partial deficit of PHP at 30 min of PhTx treatment.

We next intended to test whether the two scenarios would result in a qualitatively similar scenario in the context of MB memory formation. Thus, we used the same strategy as above described when analyzing BRP, means we post-developmentally knocked down *arl8* using a pan-neuronal MB driver (*OK107-Gal4*). We first stained the brains of flies expressing RNAi-Arl8 in adult MB lobes to check whether RNAi expression would grossly affect AZs. However, after 9 days of 29°C induction, no significant changes of BRP, RIM-BP and Syd1 were observed when compared to *Gal80^ts^;OK107/+* control flies (Fig. 5A), indicating that the gross architecture of KC synapses is not modified by Arl8 knockdown. Even after these 9 days of induction, Arl8 MB knockdown flies showed normal STM (Fig. 5B). At the same time, however, a drastic reduction of MTM (at 1 h) was measured (Fig. 5C). Again, as after BRP knockdown, decreased ASM (Fig. 5C, but not ARM as shown in Supplementary Fig. 5B) was responsible for this phenotype. Importantly, *Gal80^ts^;OK107/RNAi-Arl8* flies displayed normal memory scores when kept at 18°C (Fig. 5C). After 5 days of induction, IMAC MB lobes KD flies presented normal STM (Fig. 5D). Again, however, MTM, due to the ASM component, was attenuated (Fig. 5E, ARM was unaffected as shown in Supplementary Fig. 5C) in *Gal80^ts^;OK107/RNAi-IMAC* flies, whereas control flies kept at 18°C, as expected, displayed normal MTM.

Thus, we again find an association between a specific deficit in the stable expression of the structural and functional aspect of PHP, and the display of MTM but not STM. Interestingly, effect sizes seem to correlate between the NMJ phenotypes and the corresponding MTM deficits.

To further corroborate this relation, we finally tested Aplip1, as this protein is also important to effectively transport BRP/RIM-BP/Unc13A precursor material along the axon^32^. We previously^23^ showed that *aplip1*-null mutant shows only an attenuation (but not elimination) of PHP expression after PhTx treatment (measured in *glurIIA* mutant background). Again, similarly to *Gal80^ts^;OK107/RNAi-IMAC* flies, animals expressing RNAi-Aplip1 in adult MB lobes present normal STM (Fig. 5F) but a decrease of MTM 1 h due to a deficit of ASM 1h compared to their genetic controls (Fig. 5G, ARM was not affected as shown in Supplementary Fig. 5C). In the absence of Gal4 induction, flies expressing RNAi against Aplip1 in the adult MB lobes had normal MTM 1 h (Fig. 5G).

Importantly, all flies expressing RNAi directed against Arl8, IMAC or Aplip1 expression presented normal olfactory acuity and shocks sensitivity scores, indicating that they should have a largely undisturbed perception of the conditioning stimuli (Table 2).

**Table 2.**
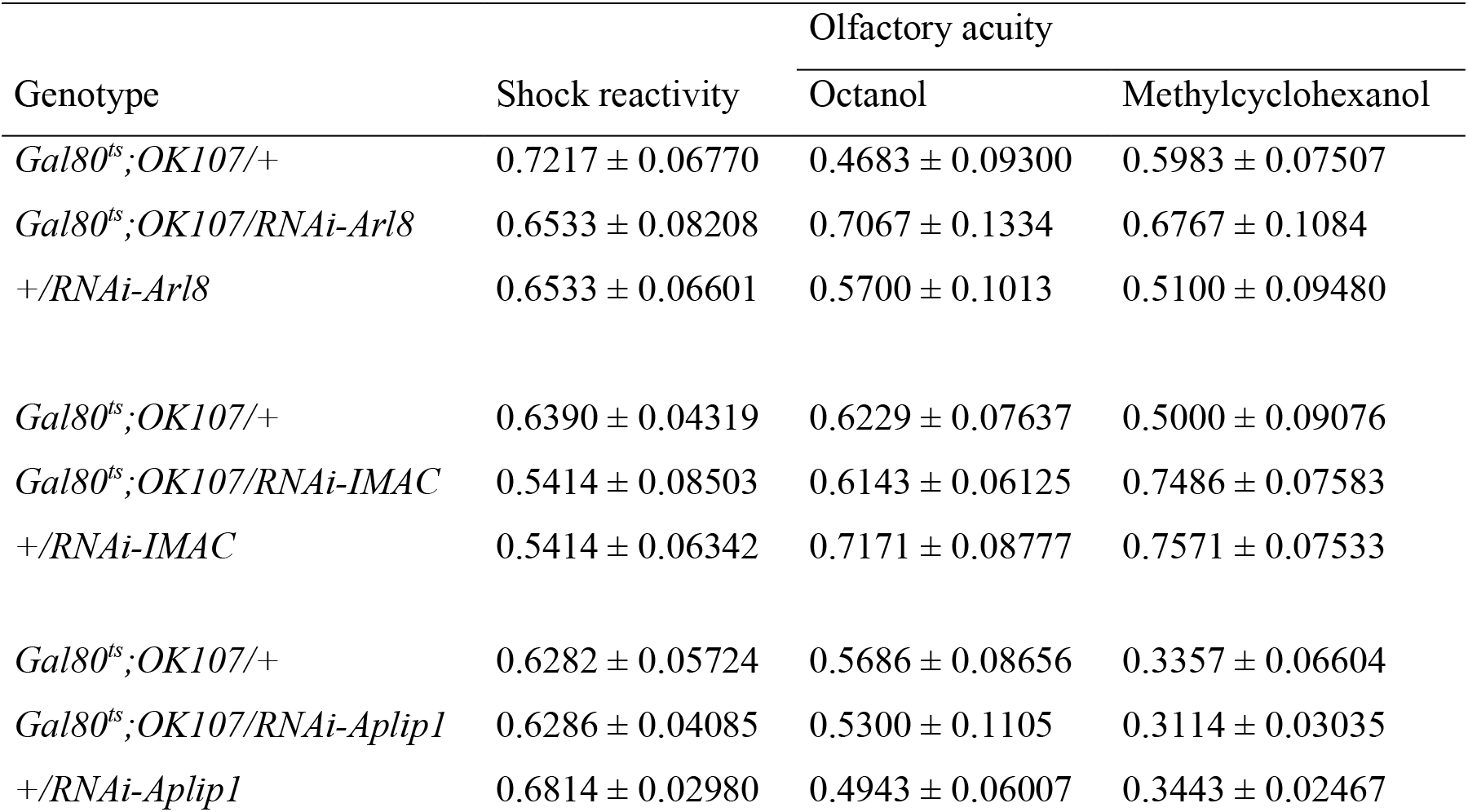
Shock reactivity and olfactory acuity of flies expressing RNAi-Arl8, -IMAC and -Aplip1 in the adult MB lobes. Data are shown as means ± SEM. After 9 d of induction, Arl8 KD flies show normal shock reactivity (*F(_3,18_)* = 0.2978, *p* = 0.7467, *n* = 6) and normal olfactory acuity for octanol (*F(_3,18_)* = 1.169, *p* = 0.3373, *n* = 6) and methycyclohexanol (*F(_3,18_)* = 0.7911, *p* = 0.4714, *n* = 6). After 5 d of induction, IMAC or Aplip1 KD flies show normal shock reactivity (RNAi-IMAC: *F(_3,24_)* = 0.7242, *p* = 0.4964, *n* ≥ 7; RNAi-Aplip1: *F(_3,25_*) = 0.3342, *p* = 0.7195, *n* ≥ 7) and normal olfactory acuity for octanol (RNAi-IMAC: *F(_3,21_)* = 0.5652, *p* = 0.5780, *n* = 7; RNAi-Aplip1: *F(_3,21_)* = 0.1776, *p* = 0.8387, *n* = 7) and methycyclohexanol (RNAi-IMAC: *F(_3,21_)* = 3.254, *p* = 0.0622, *n* = 7; RNAi-Aplip1: *F(_3,21_)* = 0.1479 *p* = 0.8635, *n* = 7).

Notably, our combined NMJ/MB analysis further supports that indeed presynaptic active zone remodeling orchestrated by the transport of BRP-containing complexes is a precondition to effectively consolidate aversive olfactory memories at one hour after conditioning. The fact that the knock-down of transport factors, not being proper active zone proteins and so not directly involved in synaptic release, resulted in this specific phenotype further supports the existence of a specific regulatory relation here.

### Homeostatic active zone plasticity via BRP is dispensable for forming aversive long-term memory

Finally, we wanted to examine a putative role of homeostatic active zone plasticity also at later points of aversive olfactory memory formation. We thus investigated whether later memory components than the so far tested 1 h MTM would similarly be affected by BRP KD. Thus, we again analyzed flies expressing RNAi-BRP^B3-C8^ in the adult MB lobes 5 days after temperature-shift triggered induction and checked them 3 h after conditioning. These *Gal80^ts^;OK107/RNAi-BRP^B3-C8^* flies also displayed a severe MTM deficit due to a loss of ASM while the ARM component (*per se* strong at 3h after conditioning) was unaffected (Fig. 6A). On contrast, control flies continuously kept at 18°C presented normal 3 h MTM (Fig. 6A) as expected. Finally, we addressed Long-Term Memory (LTM) 24 h after aversive conditioning. To generate aversive LTM, spaced training consisting of three training rounds was applied. Notably, however, LTM was obviously unaffected by BRP knockdown (Fig. 6B).

**Figure 6.**
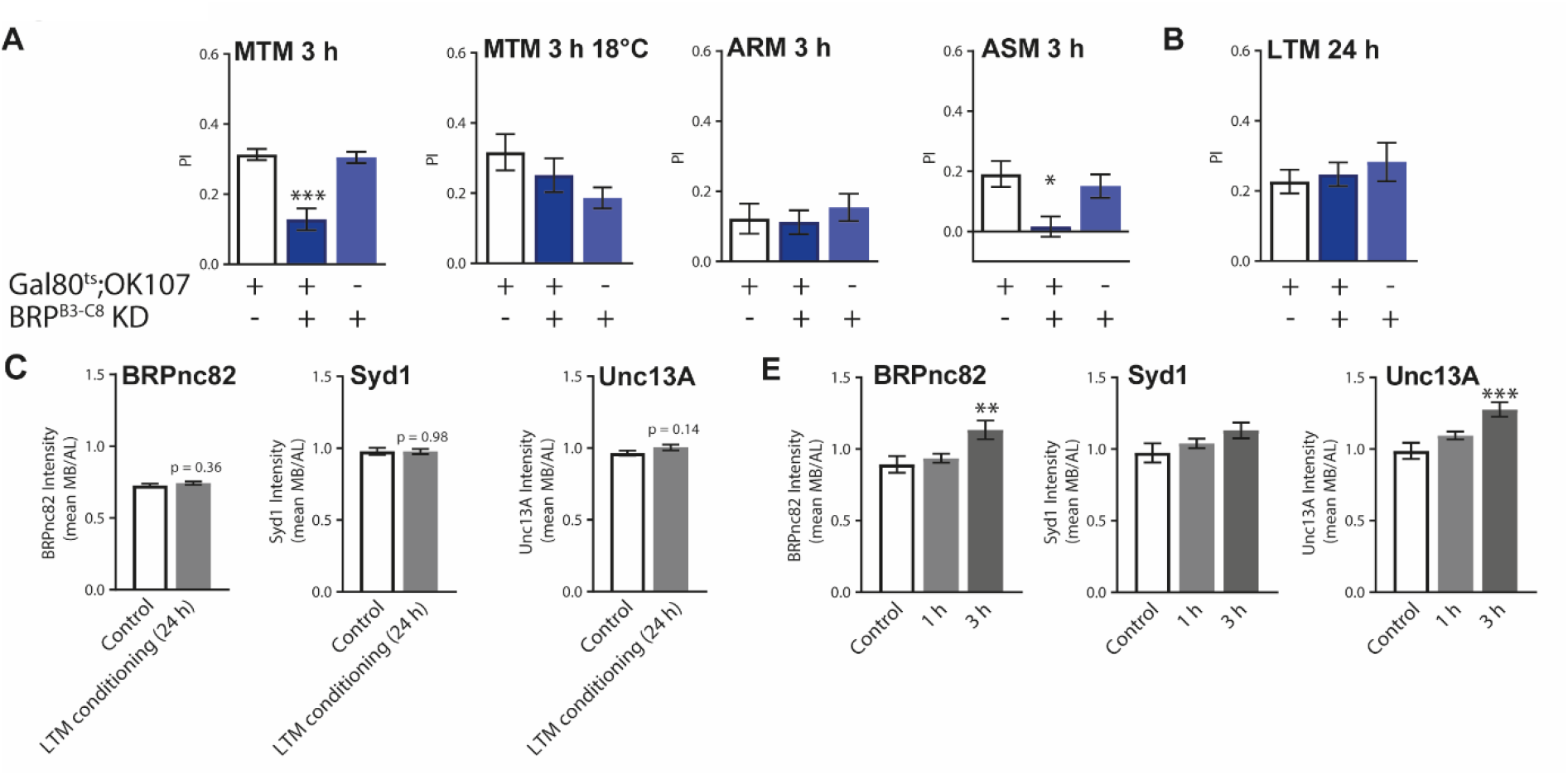
Homeostatic AZ plasticity is not essential for forming aversive LTM. A. Flies expressing RNAi-BRP^B3-C8^ in the adult MB lobes exhibit a defect of MTM 3 h (*F*(_3,45_) = 20.07, *p* < 0.0001, *n* ≥ 12; *post hoc* Tukey’s multiple comparisons test, *Gal80^ts^;OK107/+* vs *Gal80^ts^;OK107/RNAi-BRP^B3-C8^* \*\*\**p* < 0.001, *+/RNAi-BRP^B3-C8^* vs *Gal80^ts^;OK107/RNAi-BRP^B3-C8^* \*\*\**p* < 0.001), with a deficit of ASM 3 h (*F*(_3,59_) = 5.779, *p* = 0.0052, *n* ≥ 18; *post hoc* Tukey’s multiple comparisons test, *Gal80^ts^;OK107/+* vs *Gal80^ts^;OK107/RNAi-BRP^B3-C8^* \*\**p* < 0.01, *+/RNAi-BRP^B3-C8^* vs *Gal80^ts^;OK107/RNAi-BRP^B3-C8^* \**p* < 0.05), and a normal ARM 3 h (*F*(_3,59_) = 0.3281, *p* = 0.7217, *n* ≥ 18). Without induction, those flies have normal MTM 3 h (*F*(_3,18_) = 2.146, *p* = 0.1514, *n* = 6). B. After 5 d of induction, *Gal80^ts^;OK107/RNAi-BRP^B3-C8^* flies have normal LTM scores 24 h after spaced conditioning (*F*(_3,28_) = 0.4841, *p* = 0.6227, *n* ≥ 7). C. Quantification of BRP ^nc82^, Syd1 and Unc13A staining intensity in MB lobes of *w1118* flies 24 h after spaced conditioning (BRP ^nc82^: *t-test*, *p* = 0.3629, *n* = 16; Syd1: *t-test*, *p* = 0.9841, *n* = 16; Unc13A: *t-test*, *p* = 0.1447, *n* = 16). D. Quantification of BRP ^nc82^, Syd1 and Unc13A staining intensity in MB lobes of *w1118* flies 1 h and 3 h after spaced conditioning (BRP ^nc82^: *F*(_3,45_) = 5.929, *p* = 0.0054, *n* ≥ 13; *post hoc* Tukey’s multiple comparisons test, *Control* vs *3 h* \*\**p* < 0.01; Syd1: *F*(_3,45_) = 2.263, *p* = 0.1166, *n* ≥ 13; Unc13A: *F*(_3,45_) = 10.11, *p* = 0.0003, *n* ≥ 13; *post hoc* Tukey’s multiple comparisons test, *Control* vs *3 h* \*\*\**p* < 0.001).

This could mean that the decrease of BRP is not severe enough to affect LTM. Alternatively, cell types other than KCs might contribute to aversive olfactory LTM formation^35–37^. It might, however, be also possible that presynaptic scaling gets restricted to a time window of few hours only in which it would be critical to retrieve memories, while processes finally promoting LTM formation might sufficiently pertain even when presynaptic scaling is attenuated. Indeed, 24 h after spaced conditioning, MB lobes did not show changes in BRP, Syd1 or Unc13A staining intensity in comparison to untrained controls (Fig. 6C). It might be argued that space training might not result in presynaptic scaling. However, when analyzed 3 h after 3 cycles of “spaced” conditioning, BRP/Unc13A staining intensity significantly increased (Fig. 6D) similarly to the single round protocol used above.

In short, homeostatic active zone remodeling within KCs might be needed to successfully retrieve aversive memories over a few hours time window after conditioning, but neither for immediate recall nor for long-term memory display. Hence, this form of plasticity might essentially be needed to allow the circuit to successfully retrieve memories in an intermediate time window, while the memory trace seemingly successfully formed directly after training might can propagate in the absence of this homeostatic memory component.

## Discussion

Two partially antagonistic types of plasticity are supposed to be involved in stable information encoding of the brain: Hebbian plasticity, a fast positive feed-back type of plasticity leading to an increase of relevant connectivity, and homeostatic plasticity, supposed to operate via slower negative feed-back mechanisms to bring the circuit back to its rest state^4, 38^. Likely, to preserve memories, animals need to maintain specific subsets of synaptic changes, while overall synaptic strength distribution and excitation/inhibition balances must remain stable, at least on the long-term. While those partly opposite but synergistic processes probably converge on similar synaptic components and compartments, how exactly their interactions relate during initial information storage and the subsequent formation of stable memories is largely unknown.

Historically, *post*synaptic plasticity mechanisms have been extensively worked out, and processes targeting postsynaptic neurotransmitter receptors have been convincingly connected to learning and memory^5, 39^. However, the necessity of using postsynaptic neurons as “reporters” of presynaptic activity (and, thus, setup “paired recordings”) has imposed an additional obstacle specific to functionally study *pre*synaptic forms of long-term plasticity; and the cellular and molecular processes modulating transmission strength by targeting the presynaptic active zones and the associated release machinery are way less characterized^40^. Consequently, while widely expressed by excitatory and inhibitory synapses of mammalian brains^40^, the behavioral relevance of longer-term presynaptic plasticity remains largely obscure. Notably, presynaptic plasticity recently was proposed to alleviate the stability-plasticity dilemma during lifelong learning^41^. A major challenge now is to identify convergence pathways unifying the obviously distinct presynaptic plasticity components, and to connect them to their behavioral roles.

Recent work suggested that the presynaptic plastic potentiation of release which can be efficiently scrutinized at larval NMJ synapses consists of distinct components^23, 24^. This work had used *glurIIA* chronically mutant background as a surrogate for long-term presynaptic plasticity addressing components specifically needed for stable expression of PHP^42–45^. We here turned to investigate the PHP 30 minutes after PhTx application, demonstrating a direct temporal trajectory of obviously distinct components “initiating but also sustaining” the course of presynaptic plasticity. To do so, we electro-physiologically and genetically study NMJ active zones 10 and 30 minutes after they were triggered towards PHP by PhTx application. Our results clearly show that BRP, member of the generic ELKS family of AZ scaffold proteins, is essential specifically for stable expression but not for PHP induction. In contrast, RIM-BP was found essential for PHP induction “already”. We suggest that this reflects the fact that RIM-BP is directly involved in mediating SVs release site functionality via its interaction with (m)unc13 release factors, shown for Unc13A at *Drosophila*^7, 15^ and munc13-1 at mammalian synapses^46^.

Our data suggest that BRP-driven “nanoscale” remodeling of active zones can promote release site formation is likely subsequent to acute modification of already existing release sites. This is interesting also in comparison to postsynaptic plasticity at glutamatergic synapses. Here, it is thought that the plasticity trajectory starts via the acute modulation of glutamate receptors via phosphorylation, only later followed by physical incorporation of additional glutamate receptors in the range of tens of minutes^47^. Notably, at rodent hippocampal mossy fiber synapses, electrophysiological and ultrastructural analysis demonstrate changes in release probability of existing release sites to be followed by the functional addition of more release sites^40^. Thus, it will be very interesting to reconcile the presynaptic plasticity sequence between rodents and *Drosophila*. In this regard, the *Drosophila* NMJ synapse with its unique genetic and physiological accessibility started to allow for a dissection of the molecular/cellular mechanisms and trajectory of presynaptic plasticity^11, 23, 24^, and this manuscript adds to the notion that active zone remodeling is indeed a multi-step process.

Notably, we found that all those AZ-related components whose elimination provoked a specific deficit in stable expression (but not induction) of NMJ PHP were also specifically needed for consolidation (1 h) of aversive memories but at the same time dispensable for aversive short-term memory formation in adult MB KCs. Indeed, we observed a correlation between the severity of the PHP expression deficit and the effect size concerning MTM attenuation (e.g. compare PHP 30 min and MTM scores for arl8 in Fig. 4C and Fig. 5C, with IMAC in Fig. 4F and Fig. 5E respectively). Given that several molecular factors, including transport proteins not directly physically associated with AZs, fulfilled this correlation, it appears most likely that indeed, it is this kind of active zone remodeling process that consolidates memories at MB AZs. That we find the induction process for both memory formation and NMJ PHP to continue apparently undisturbed while the later expression process was severely affected is arguing in direction of a truly specific functional relation and should rebut arguments such as base-line synaptic defects being responsible here (also note that we achieved behavioral phenotypes by comparatively “mild” and strictly post-developmental knockdowns). That indeed we face a *plasticity sequence operating along time at a coherent set of synapses* is a suggestive possibility, which, however, clearly needs additional analysis. The fact that aversive STM and MTM in experiments acutely blocking synaptic release (via *shibire^ts^)* was predominantly mapped to either γ- or α/β neurons is not in immediate match with this possibility^18, 48, 49^.

However, the direct imaging-based demonstration of a conditioning-dependent upscaling of BRP and Unc13A (^22^; this study Fig. 2A) observed in KCs and over the MB lobes obviously matches with our finding that the homeostatic active zone directed machinery (needed for BRP/Unc13A scaling) is of specific importance for memory consolidation or expression.

Surprisingly, we found that a knock-down of BRP in the adult MB lobes did not affect LTM whereas MTM is decreased both at 1 h and at 3 h. Such a phenotype, deficit of MTM but subsequent memory phases being intact, was only rarely observed before (*Nep2*-RNAi in adult DPM neurons^37^, *synapsin* mutants with memory deficits at 3 min and 1 h but normal memory at 5 h^50^). This on one hand would reinforce the idea that MTM and LTM are formed by either separate AZs circuits or on the other hand required different set of proteins in the same lobes. However, it also might be that presynaptic scaling is needed only during a few hours time window, and that the “trace” leading to LTM is either mediated by independent molecular/synaptic mechanisms or distinct circuits.

What might be the circuit level role of presynaptic scaling? Notably, the acute formation of aversive short-term memory formation was shown to trigger synaptic depression at the KC::MBON synapse in the respective MB compartment^51^ ; thus, we arrive at the interesting possibility that the synaptic scaling process might indeed follow in time the initial depression (generating STM traces) to homeostatically refine the circuits in direction of equilibrated synaptic weights and excitability, in effect preparing for subsequent rounds of learning. Indeed, networks are meant to operate close to “criticality” by using distinct scale plasticity mechanisms^52, 53^. Our findings now provide a direct experimental angle to investigate this hypothesis behaviorally, genetically and physiologically.

Interestingly, dopamine and NO as co-transmitters from defined sets of dopaminergic neurons were recently shown to within KCs trigger memories of opposing valence, where in comparison to the dopamine effect, the NO-dependent effect develops slowly, requires longer training than dopamine-dependent memory, and shortens memory retention^54^. While in this scenario two subsequent antagonistic, versus in our story two subsequent synergistic memory components, are generated, the scheme of two molecular-mechanistically separated presynaptic memory components following each other is intriguing. Further exploring the exact cellular and molecular mechanisms coupling these phenomena across the plasticity sequence might well be warranting.

Notably, it previously has been described that constitutive knock-down of BRP in the MB lobes (also using *OK107-Gal4* but constitutive without the use of Gal80^ts^) leads to an ARM deficit^55^. We wanted to see whether we would reproduce this result using another more specific pan-MB driver line, VT30559^56^. When BRP was knock-down in the MB during both development and adult stage driving expression with VT30559, flies presented a deficit of STM, normal MTM 1 h and a defect of MTM 3 h, which, however, was due to an extreme reduction of ARM (Supplementary Fig. 6A, for olfactory and shocks sensitivity controls see Supplementary Table). Indeed, the ASM component was not affected (Supplementary Fig. 6A). The discrepancy between results obtained after knock-down of BRP either during development and adulthood or only in adulthood is likely due to the capacity of the brain to adapt after long time BRP deficit during development in regard to the ASM component, whereas the presence of the ARM 3 h defect after constitutive KD probably reflects a structural deficit due to prolonged absence of BRP in the circuits needed for ARM formation. This “swapping of requirements” warrants future scrutiny.

RIM-BP is the only protein linked to synaptic vesicles release that we tested and the only protein we found involved in STM. We finally wanted to verify whether this role of RIM-BP in early memory is specifically caused by the fact that RIM-BP is involved in synaptic release or if the deficit observed is a consequence of an impairment of the AZ structure due to the absence of one of its linkers to synaptic vesicles. We thus decided to test another protein linked to synaptic vesicles but not to the AZ itself. We chose a SNARE protein attached to the vesicles, involved in the docking and release of SVs: Synaptotagmin1 (Syt1)^57–59^. Similarly to RIM-BP KD, Syt1 KD in adult MB lobes lead to deficit of STM and MTM 1 h (Supplementary Fig. 6B for STM and 6C for MTM 1 h, see olfaction and shocks sensitivity controls in Supplementary Table) indicating that altering the synaptic vesicles release itself leads to STM defect.

Importantly, presynaptic homeostasis was just shown to oppose the disease progression in a mouse model of ALS-Like Degeneration^60^. Thus, deciphering the exact mechanistic role of presynaptic homeostatic plasticity and its obviously distinct mechanistic components might be equally relevant for basic but also medically oriented research.

## Materials and methods

### Drosophila stocks

*Drosophila* wild-type strain *w1118* and mutant flies were raised on conventional cornmeal-agar medium in 60 % humidity in a 12 h light/dark cycle at 25°C for experiments on larva and 18°C for experiments in adults. All stains used for memory experiments were outcrossed to the *w1118* background. *brp*-null (*brp^Δ6^*^.1^/*brp^69^* ^23^) and *arl8*-null (PBac(RB)Giee00336^30^, Bloomington 17846) and *imac*-hypomorphe *unc-104^bris^*/*unc-104^d11024^* (*unc-104^bris^* ^61^, *unc-104^d11024^* Bloomington 19346) mutants flies were used. RNAi stocks were obtained from the Vienna Drosophila Resource Center (Austria): RNAi-Arl8 (VDRC 26085), RNAi-Aplip1 (VDRC 50007), RNAi-IMAC (VDRC 23465) and RNAi-Syt1 (VDRC 8874). RNAi-BRP^B3-C8^ has been described in Wagh et al.^27^. RNAi-RIM-BP flies have been obtained after design of the RNAi sequence by our laboratory (Forward: 5’-CTAGCAGTGGGCACCGACAATCAGCCACCTAGTTATATTCAAGCATAGGTGGCTGATTGTCGGTGCCCGCG-3’; Reverse: 5’-AATTC GCGGGCACCGACAATCAGCCACCTATGCTTGAATATAACTAGGTGGCTGATTGTGGTGCCCACTG-3’) and injection by BestGene Inc. The *tubulin-Gal80^ts^;OK107* driver (*Gal80^ts^;OK107*) was used for conditional expression in the MB, *tubulin-Gal80^ts^;c739* (*Gal80^ts^;c739*) for conditional expression in α/β neurons, and *VT30559* (v206077; Vienna Drosophila Resource-Center, Austria) for constitutive expression in the MB. To induce RNAi expression specifically in adults, the TARGET system was used as described by McGuire et al.^26^: flies were kept for 5 days at 29°C before staining, conditioning and until memory test for LTM analysis for all RNAi expressing flies, except flies expressing RNAi-Arl8 and flies expressing RNAi-BRP^B3-C8^ in one STM experiment which were kept 9 days.

### Behavior: olfactory associative aversive conditioning

Flies were trained using the classical olfactory aversive conditioning protocols described by Tully & Quinn^62^. Training and testing were performed in climate-controlled boxes at 25°C in 80 % humidity under dim red light. At 2-3 days old, flies were transferred to fresh food vials and either put at 29°C for RNAi induction or stayed at 18°C for the non-induced controls. Conditioning was performed on groups of around 40-50 flies with 3-octanol (around 95 % purity; Sigma-Aldrich) and 4-methylcyclohexanol (99 % purity; Sigma-Aldrich). Odors were diluted at 1:100 in paraffin oil and presented in 14 mm cups. A current of 120 AC was used as a behavioral reinforcer. Memory conditioning and tests were performed with a T-maze apparatus^62^. In a single-cycle training, groups of flies were presented with one odor (CS^+^) paired with electrical shock (US; 12 times for one minute). After one minute of pure air-flow, the second odor (CS^-^) was presented without the shock for another minute. During the test phase, flies were given 1 min to choose between 2 arms, giving each a distinct odor. An index was calculated as the difference between the numbers of flies in each arm divided by the sum of flies in both arms. The average of two reciprocal experiments gave a performance index (PI). The values of PI ranges from 0 to 1, where 0 means no learning (50:50 distribution of flies) and a value of 1 means complete learning (all flies avoided the conditioned odor). To analyze LTM, flies were kept after training in standard food vials at 29°C for 24 h until memory tests were performed. For olfactory acuity and shock reactivity, around 50 flies were put in a choice position between either one odor and air for one minute or electric shocks and no-shocks respectively.

### Immunostaining

Brains were dissected in ice-cold hemolymph-like saline (HL3; composition in mM: NaCl 70, KCl 5, MgCl2 20, NaHCO3 10, trehalose 5, sucrose 115, HEPES 5, pH adjusted to 7.2) solution and immediately fixed in 4 % paraformaldehyde (PFA) for 30 min at room temperature under stirring. Samples were incubated for 2 h in phosphate-buffered saline (PBS) containing 1 % Triton X-100 (PBT) containing 10 % normal goat serum (NGS). Subsequently, the samples were incubated in the primary antibody solution diluted in PBT-5 % NGS at 4°C under stirring for 48 h. Samples were washed six times for at least 30 min each in PBT at room temperature, and subsequently incubated with secondary antibody solution diluted in PBT-5 % NGS at 4°C overnight. Brains were washed at room temperature six times for at least 30 min each in PBT. Finally, the mounting was done in Vectashield (Vector Laboratories) on glass slides. The following primary antibodies were used: mouse BRP (BRP^nc82^, DSHB; diluted 1:50), rabbit RIM-BP^C-term 63^ (diluted 1:250), rabbit Syd1^64^ (diluted 1:250), guinea pig Unc13A^15^ (diluted 1:500). The following secondary antibodies were used: Alexa Fluor 488-coupled goat anti-mouse (Invitrogen A11029; diluted 1:500), Cy-3-coupled goat anti-rabbit (Biozol 111-165-006; diluted 1:500), Cy-5-coupled goat anti-rabbit (ThermoFisher A-10523; diluted 1:500), Cy-3-coupled goat anti-guinea pig (Abcam ab102370; diluted 1:500).

Larvae were dissected and stained as previously described^23^ with BRP (mouse BRP^nc82^, DSHB; diluted 1:200) and secondary Alexa Fluor 488-coupled goat anti-mouse (Invitrogen Cat#A-11001; RRID: AB_2534069; diluted 1:500). The glutamate receptor blocker Philanthotoxin-433 (PhTx, Aobious, MA, USA) was prepared as a 4 mM stock solution in dH_2_0. Rapid homeostatic plasticity was induced through pharmacological challenge with 50µM PhTx in calcium-free HL3 at room temperature. Controls were similarly treated by substituting PhTx with dH_2_0. Briefly, the larvae were immobilized with insect pins on a rubber dissection pad, cut open dorsally between the dorsal tracheal trunks, avoiding excessive stretching or tissue damage. The semi-intact larvae were incubated with PhTx for 10 min. The preparation was completed by flattening the body wall using insect pins to expose the muscles. The tissue was fixed with ice-cold 4 % PFA in 0.1 mM PBS for 10 min and all extraneous tissue was removed. Larvae were then processed for immunohistochemistry and mounted in Vectashield (Vector Labs, CA, USA).

### Confocal imaging and data processing

Adult brain samples were imaged using Leica SP8 confocal microscope equipped with x20 apochromat oil-immersion Leica objective (NA=0.75) and x40 apochromat oil-immersion objective (NA=1.30). Alexa Fluor 488 was excited at 488 nm, Cy3 at 561 nm and Cy5 at 633 nm wavelengths. Samples were scanned using LAS X software (3.5.2.18963) at 0.5 µm sections in the z direction. All images were acquired at 8-bits grayscale. Segmentation of the image stacks were processed using the Amira® software (Visage Imaging GmbH). The first step was to define a unique label for each region in the first (BRP^nc82^) fluorescent channel for the RNAi experiment and the second (Syd1) fluorescent channel for the experiment after conditioning. A full statistical analysis of the image data associated with the segmented materials was obtained by applying the Material Statistics module of the Amira software, in which the mean gray value of the interior region is calculated. To avoid difference of global staining between different groups, the intensity of the staining of the MB lobes and calyx was normalized on the intensity of Antennal Lobes (AL) and Protocerebral Bridge (PB) respectively for quantification purpose. The median voxel values of the regions were compared, as measured in individual adult brains, in order to evaluate the synaptic marker label.

For larval NMJ, confocal microscopy was performed with a Leica TCS SP8 inverted confocal microscope (Leica DMI 6000, Leica Microsystems, Germany). All images were acquired at room temperature using LCS AF software. (Leica Microsystems, Germany). Confocal imaging was performed using a 63×1.4 NA oil immersion objective. Images of abdominal muscle 4 Type-1b NMJs were obtained from fixed larval preparations for all experiments. Images were acquired in line scanning mode with a pixel size 75.16nm*75.16nm and with a z-step of 0.25 µm.

All brain and NMJ representative images were processed using the ImageJ/Fiji software (1.52P, https://imagej.net/software/fiji/) for adjusting brightness with the brightness/contrast function. Images shown in a comparative figure were processed with exactly the same parameters.

### Electrophysiology

Electrophysiological recordings were performed at room temperature on muscle 6 of 3^rd^ instar larval NMJs in the abdominal segments A2 and A3. The larvae were incubated in Ca^2+^-free modified HL3 solution (composition (in mM): NaCl 70, KCl 5, MgCl_2_ 10, NaHCO_3_ 10, trehalose 5, sucrose 115, HEPES 5, pH adjusted to 7.2) containing either 50 µM PhTx (or an equal volume of H2O for controls) for the indicated amount of time (10 min or 30 min) and dissected toward the end of the incubation time (∼2 min). Larvae were then washed 3 times in modified HL3 without PhTx and recordings were obtained in a bath solution of modified HL3 containing 0.4 mM CaCl_2_. Glass electrodes were pulled using a Flaming Brown Model P-97 micropipette puller (Sutter Instrument, CA, USA). Recordings were made using an Axoclamp 2 B amplifier with HS-2A x0.1 headstage (Molecular Devices, CA, USA) and on a BX51WI Olympus microscope with a 40X LUMPlanFL/IR water immersion objective (Olympus Corporation, Shinjuku, Tokyo, Japan). mEPSPs were recorded for 90 seconds. eEPSPs were recorded after stimulating the appropriate motoneuron bundle with 8 V, 300 µs at 0.2 Hz using an S48 Stimulator (Grass Instruments, Astro-Med, Inc., RI, USA). Signals were digitized at 10 kHz using an Axon Digidata 1322 A digitizer (Molecular Devices, CA, USA) and low pass filtered at 1 kHz using an LPBF-48DG output filter (NPI Electronic, Tamm, Germany). The recordings were analysed with pClamp 10 (Molecular Devices, Sunnyvale, CA, USA), Graphpad Prism 6 (Graphpad Software, Inc., San Diego, CA, USA) and MATLAB R2010b (Mathworks, Natick, MA, USA). Stimulation artifacts of eEPSPs were removed for clarity.

Current clamp recordings were performed as previously described^65^. Recordings were made from cells with an initial Vm between –40 and –70 mV, and input resistances of ≥ 4 MΩ, using intracellular electrodes with resistances of 30-60 MΩ, filled with 3 M KCl. mEPSPs were further filtered with a 500 Hz Gaussian low-pass filter. Using a single template for all cells, mEPSPs were identified and analysed, noting the mean mEPSP amplitude per cell. An average trace was generated from 20 eEPSP traces per cell. The amplitude of the average eEPSP trace was divided by the mean mEPSP amplitude, for each respective cell, to determine the quantal content.

### Statistical analysis

Data was analyzed using Prism (GraphPad Software, CA, USA).

For adult brain staining experiments, differences among multiple groups were tested by one-way ANOVA with Tukey’s post hoc test, whereas differences between two groups were test by t-test.

Memory scores are displayed as mean ± SEM. For behavioral experiments, scores resulting from all genotypes were analyzed using one-way ANOVA followed, if significant at p ≤ 0.05, by Tukey’s multiple-comparisons tests. For memory experiments, the overall ANOVA p-value is given in the legends, along with the value of the corresponding Fisher distribution F(x,y), where x is the number of degrees of freedom for groups and y is the total number of degrees of freedom for the distribution. Asterisks on the figure denote the least significant of the pairwise post hoc comparisons between the genotype of interest and its controls following the usual nomenclature (ns, (not significant) p > 0.05; *, p ≤ 0.05; **, p ≤ 0.01; ***, p ≤ 0.001). The number of independent experiments (n) are mentioned in the figure legends.

For larval NMJ experiments, non-parametric t-test or Mann-Whitney U test was used for datasets with two groups. For datasets with three or more groups, non-parametric Kruskal-Wallis test followed by Dunn’s multiple comparison test or one-way ANOVA followed by Tukey’s multiple comparison test was used. For all datasets failing a D’Agostino & Pearson normality test and all immunostaining data, Mann-Whitney U test or non-parametric Kruskal-Wallis test followed by Dunn’s multiple comparison test was used. Statistical parameters are stated in the figure legends. Data is represented as mean ± SEM. Statistical significance is denoted in the graphs as asterisks: *, p < 0.05; **, p < 0.01; ***, p < 0.001; ns, (not significant), p > 0.05.

## Author Contributions

O.T. and S.J.S. designed the experiments and wrote the paper. O.T. performed fly husbandry and maintenance, memory experiments, adult confocal microscopy and analyzed those data.

N.R. and M.J.F.E. performed electrophysiological experiments, NMJ microscopy and analyzed those data.

## Acknowledgements

This work was supported by grants to S.J.S. (SFB1315 A08) and O.T. (Fondation Fyssen, Bettencourt Schueller Foundation). We want to thank Carla Hamildago for help on Syt1 experiments.

---

**Supplementary figure 3.**
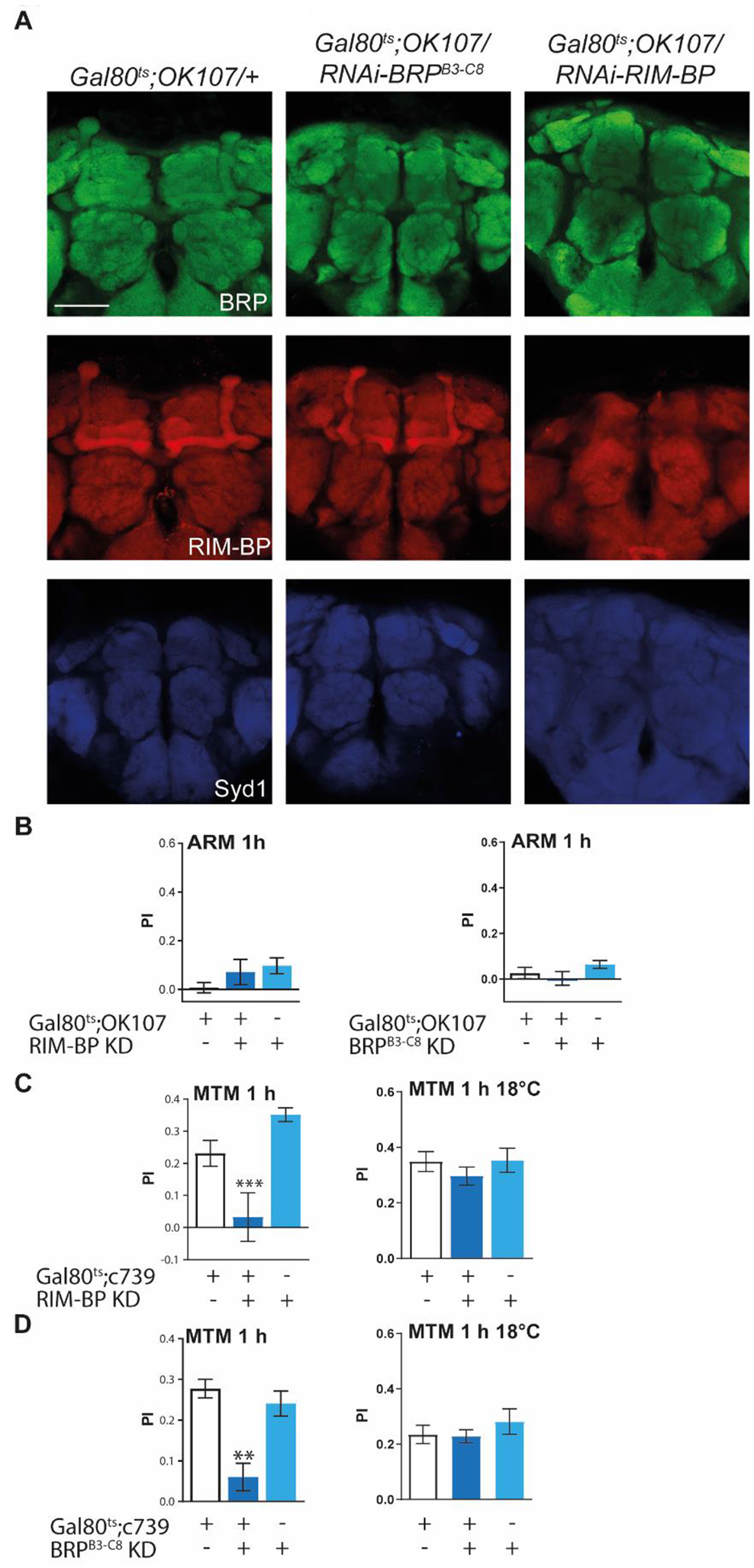
A. Representative confocal images for the staining quantification presented in Figure3A. Scale bar: 50 µm. B. Flies expressing RNAi-RIM-BP (*F*(_3,37_) = 1.439, *p* = 0.2513, *n* ≥ 12) and RNAi-BRP^B3-C8^ (*F*(_3,60_) = 1.635, *p* = 0.2040, *n* = 20) have normal ARM 1 h. C. Flies expressing RNAi-RIM-BP in the adult α/β MB lobes present MTM 1 h deficit (*F*(_3,24_) = 10.02, *p* = 0.0009, *n* = 8; *post hoc* Tukey’s multiple comparisons test, *Gal80^ts^;c739/+* vs *Gal80^ts^;c739/RNAi-RIM-BP* \**p* < 0.05, +/*RNAi-RIM-BP* vs *Gal80^ts^;c739/RNAi-RIM-BP* ***p < 0.001). Without induction, flies have normal MTM 1 h (*F*(_3,27_) = 0.7129, *p* = 0.5003, *n* ≥ 8). D. After induction, *Gal80^ts^;c739/RNAi-BRP^B3-C8^* flies present MTM 1 h deficit (*F*(_3,34_) = 10.87, *p* = 0.0003, *n* ≥ 19; *post hoc* Tukey’s multiple comparisons test, *Gal80^ts^;c739/+* vs *Gal80^ts^;c739/RNAi-BRP^B3-C8^* \**p* < 0.05, *+/RNAi-BRP^B3-C8^* vs *Gal80^ts^;c739/RNAi-BRP^B3-C8^* \*\*\**p* < 0.001). After 5 d at 18°C, those flies have normal MTM 1 h (*F*(_3,23_) = 0.6716, *p* = 0.5220, *n* ≥ 7).

**Supplementary figure 5.**
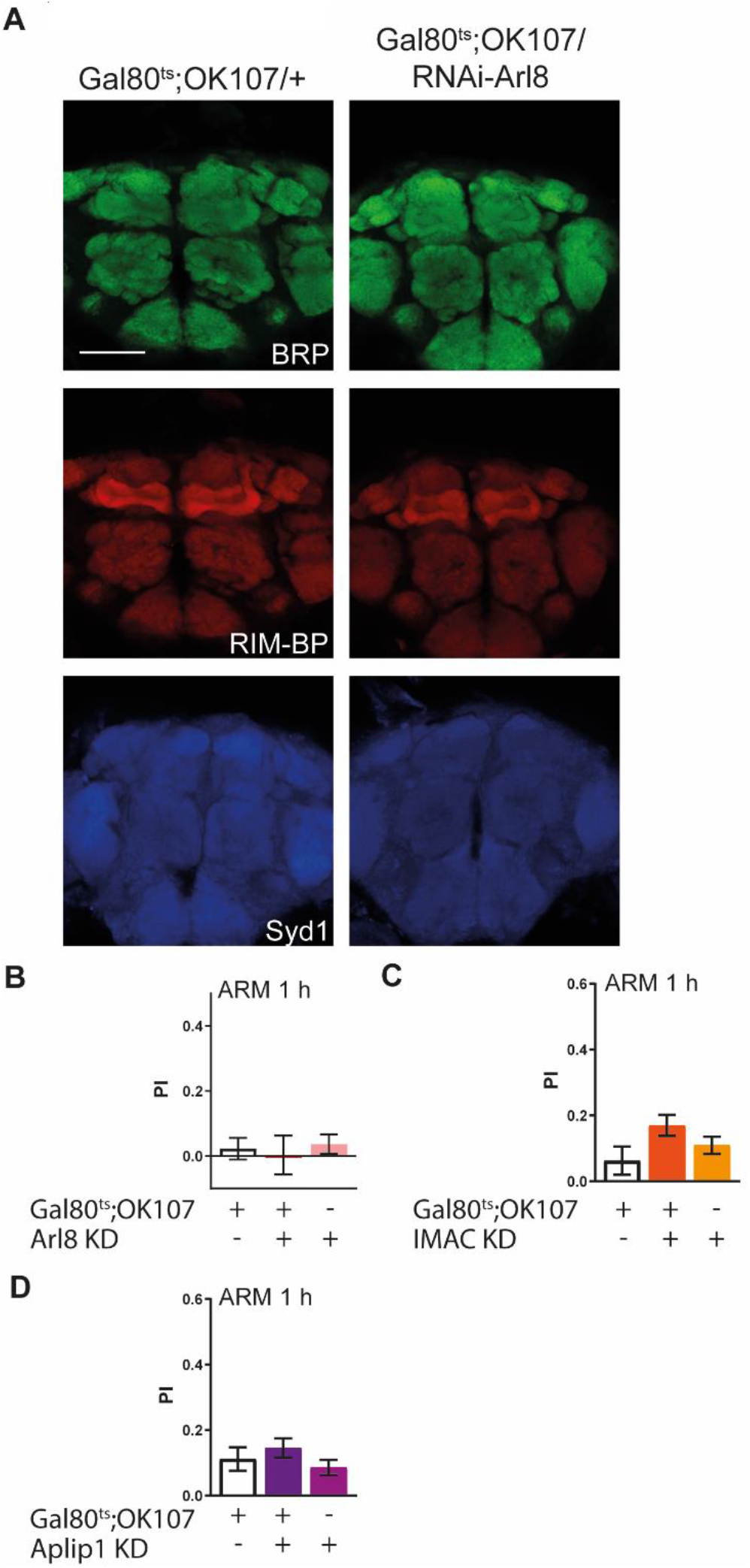
A. Representative images for the staining quantification presented in Figure 5A. Scale bar: 50 µm. B. Flies expressing RNAi-Arl8 in the adult MB lobes have normal ARM 1 h (*F*(_3,24_) = 0.1703, *p* = 0.8446, *n* ≥ 7). C. Flies expressing RNAi-IMAC in the adult MB lobes have normal ARM 1 h (*F*(_3,51_) = 2.417, *p* = 0.1000, *n* = 17). D. Flies expressing RNAi-Aplip1 in the adult MB lobes have normal ARM 1 h (*F*(_3,52_) = 0.9911, *p* = 0.3785, *n* ≥ 17).

**Supplementary Figure 6.**
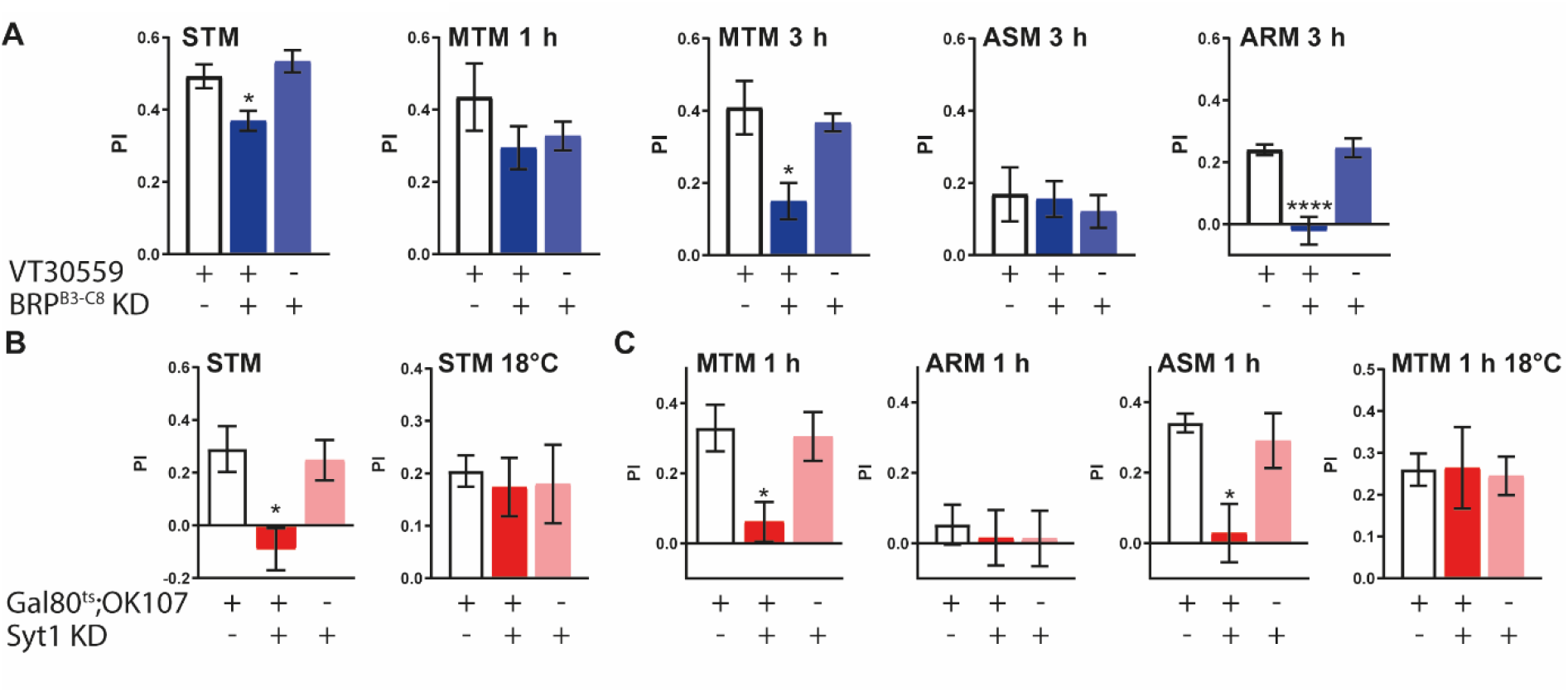
Constitutive knock-down of BRP leads to STM and 3 h ARM deficits and Synaptotagmin 1 is necessary for both STM and MTM. A. Flies expressing RNAi-BRP^B3-C8^ during development and adulthood present a defect of STM (*F*(_3,61_) = 7.667, *p* = 0.0011, *n* ≥ 20; *post hoc* Tukey’s multiple comparisons test, *VT30559/+* vs *VT30559/RNAi-BRP^B3-C8^* \**p* < 0.05, *+/RNAi-BRP^B3-C8^* vs *VT30559/RNAi-BRP^B3-C8^* \*\**p* < 0.01), have normal MTM 1 h (*F*(_3,22_) = 1.233, *p* = 0.3138, *n* ≥ 6) and a strong defect of MTM 3 h (*F*(_3,23_) = 7.425, *p* = 0.0039, *n* ≥ 6; *post hoc* Tukey’s multiple comparisons test, *VT30559/+* vs *VT30559/RNAi-BRP^B3-C8^* \*\**p* < 0.01, *+/RNAi-BRP^B3-C8^* vs *VT30559/RNAi-BRP^B3-C8^* \**p* < 0.05), with normal ASM 3 h (*F*(_3,22_) = 0.1909, *p* = 0.8278, *n* ≥ 7), and a strong deficit of ARM 3 h (*F*(_3,22_) = 21.53, *p* < 0.0001, *n* ≥ 7; *post hoc* Tukey’s multiple comparisons test, *VT30559/+* vs *VT30559/RNAi-BRP^B3-C8^* \*\*\**p* < 0.001, *+/RNAi-BRP^B3-C8^* vs *VT30559/RNAi-BRP^B3-C8^* \*\*\**p* < 0.001). B. Flies expressing RNAi-Syt1 in the adult MB lobes exhibit a defect of STM (*F(_3,36_)* = 6.209, *p* = 0.0051, *n* ≥ 11; *post hoc* Tukey’s multiple comparisons test, *Gal80^ts^;OK107/+* vs *Gal80^ts^;OK107/RNAi-Syt1* \*\**p* < 0.01, *+/RNAi-Syt1* vs *Gal80^ts^;OK107/RNAi-Syt1* \**p* < 0.05). Without induction, those flies have normal STM (*F(_3,26_)* = 0.08225, *p* = 0.9213, *n* ≥ 8). C. *Gal80^ts^;OK107/RNAi-Syt1* flies present a deficit of MTM 1 h (*F(_3,42_)* = 6.041, *p* = 0.0052, *n* ≥ 10; *post hoc* Tukey’s multiple comparisons test, *Gal80^ts^;OK107/+* vs *Gal80^ts^;OK107/RNAi-Syt1* \*\**p* < 0.01, *Gal80^ts^;OK107/RNAi-Syt1* vs *+/RNAi-Syt1* \**p* < 0.05) with normal ARM (*F(_3,25_)* = 0.1087, *p* = 0.8975, *n* ≥ 7) and a decreased ASM (*F(_2,24_)* = 6.527, *p* = 0.0062, *n* ≥ 7; *post hoc* Tukey’s multiple comparisons test, *Gal80^ts^;OK107/+* vs *Gal80^ts^;OK107/RNAi-Syt1* \*\**p* < 0.01, *Gal80^ts^;OK107/RNAi-Syt1* vs *+/RNAi-Syt1* \**p* < 0.05). Without induction, those flies present normal MTM 1 h (*F(_3,26_)* = 0.08225, *p* = 0.9213, *n* ≥ 8).

**Supplementary table.**
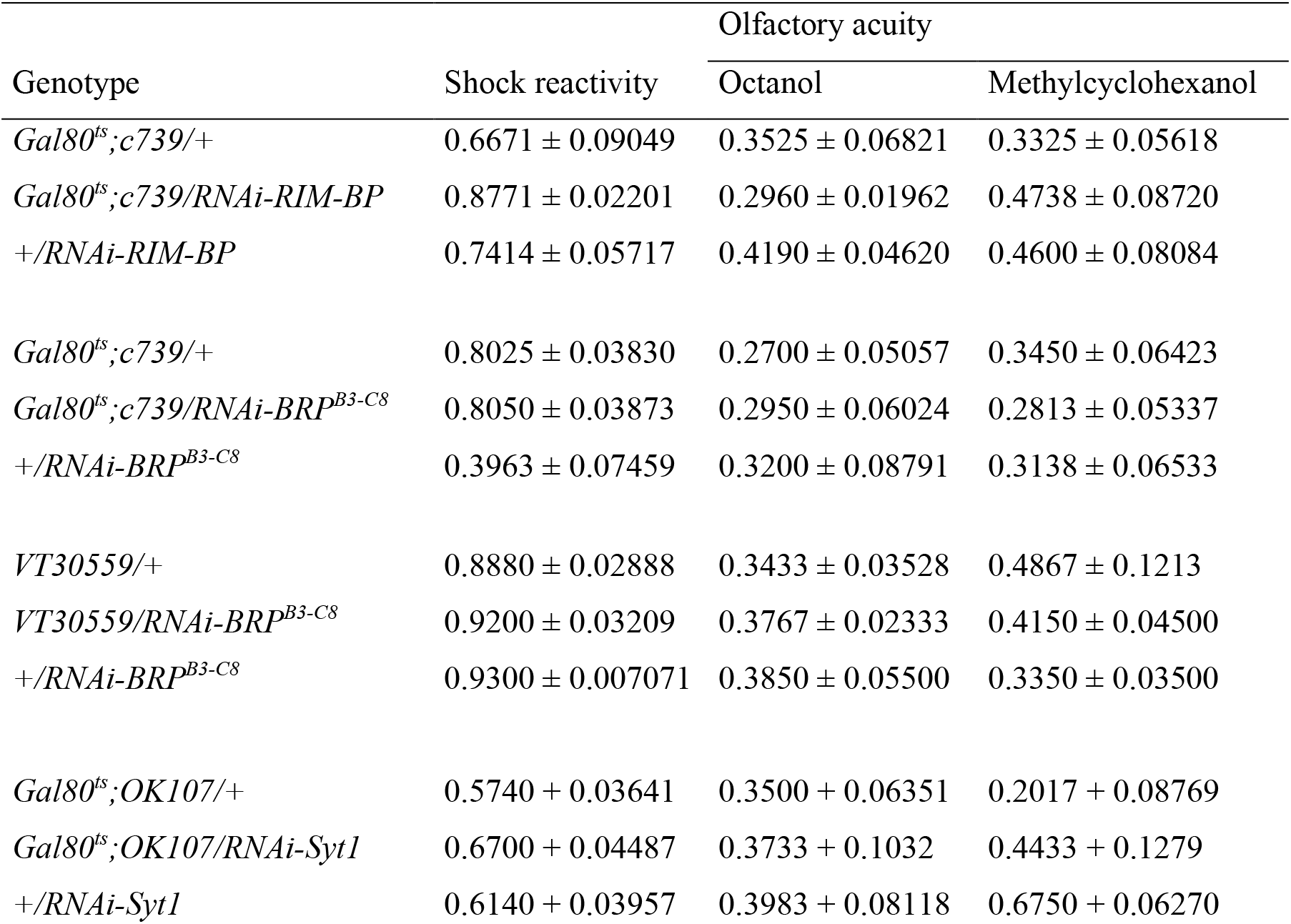
Shock reactivity and olfactory acuity of flies expressing RNAi-RIM-BP and RNAi-BRP^B3-C8^ in the adult α/β lobes, of flies expressing RNAi-BRP^B3-C8^ during development and adulthood in the MB lobes and of flies expressing RNAi-Syt1 in the adult MB lobes. Data are shown as means ± SEM. After 5 d of induction, flies expressing RNAi-RIM-BP and RNAi-BRP^B3-C8^ in the α/β lobes show normal shock reactivity (RNAi-RIM-BP: *F(_3,21_)* = 2.849, *p* = 0.0842, *n* = 7; RNAi-BRP^B3-C8^: *F(_3,24_)* = 19.47, *p* < 0.0001, *n* = 8; *post hoc* Tukey’s multiple comparisons test, *Gal80^ts^;c739/+* vs *+/RNAi-BRP^B3-C8^* \*\*\**p* < 0.001, *+/RNAi-BRP^B3-C8^* vs *Gal80^ts^;c739/RNAi-BRP^B3-C8^* \*\*\**p* < 0.001) and normal olfactory acuity for octanol (RNAi-RIM-BP: *F(_3,28_)* = 1.944, *p* = 0.1641, *n* ≥ 8; RNAi-BRP^B3-C8^: *F(_3,24_)* = 0.1348, *p* = 0.8747, *n* = 8) and methycyclohexanol (RNAi-RIM-BP: *F(_3,24_)* = 1.052, *p* = 0.3669, *n* = 8; RNAi-BRP^B3-C8^: *F(_3,24_)* = 0.2712, *p* = 0.7651, *n* = 8). Flies expressing RNAi-BRP^B3-C8^ during development and adulthood in the MB lobes show normal shock reactivity (*F(_3,18_)* = 0.3065, *p* = 0.7405, *n* = 6) and normal olfactory acuity for octanol (*F(_3,19_)* = 0.3152, *p* = 0.7341, *n* ≥ 6) and methycyclohexanol (*F(_3,19_)* = 0.02925, *p* = 0.9712, *n* ≥ 6). Flies expressing RNAi-Syt1 in the adult MB lobes show normal shock reactivity (*F(_3,15_)* = 1.411, *p* = 0.2790, *n* ≥ 5) and normal olfactory acuity for octanol (*F(_3,17_)* = 0.08237, *p* = 0.9213, *n* = 6) and methylcyclohexanol (*F(_3,17_)* = 0.2431, *p* = 0.2431, *n* = 6).

| Genotype | Shock reactivity | Olfactory acuity |  |
| --- | --- | --- | --- |
|  |  | Octanol | Methylcyclohexanol |
| *Gal80<sup>ts</sup>;c739/+* | 0.6671 ± 0.09049 | 0.3525 ± 0.06821 | 0.3325 ± 0.05618 |
| *Gal80<sup>ts</sup>;c739/RNAi-RIM-BP* | 0.8771 ± 0.02201 | 0.2960 ± 0.01962 | 0.4738 ± 0.08720 |
| *+ /RNAi-RIM-BP* | 0.7414 ± 0.05717 | 0.4190 ± 0.04620 | 0.4600 ± 0.08084 |
| *Gal80<sup>ts</sup>;c739/+* | 0.8025 ± 0.03830 | 0.2700 ± 0.05057 | 0.3450 ± 0.06423 |
| *Gal80<sup>ts</sup>;c739/RNAi-BRP<sup>B3-C8</sup>* | 0.8050 ± 0.03873 | 0.2950 ± 0.06024 | 0.2813 ± 0.05337 |
| *+ /RNAi-BRP<sup>B3-C8</sup>* | 0.3963 ± 0.07459 | 0.3200 ± 0.08791 | 0.3138 ± 0.06533 |
| *VT30559/+* | 0.8880 ± 0.02888 | 0.3433 ± 0.03528 | 0.4867 ± 0.1213 |
| *VT30559/RNAi-BRP<sup>B3-C8</sup>* | 0.9200 ± 0.03209 | 0.3767 ± 0.02333 | 0.4150 ± 0.04500 |
| *+ /RNAi-BRP<sup>B3-C8</sup>* | 0.9300 ± 0.007071 | 0.3850 ± 0.05500 | 0.3350 ± 0.03500 |
| *Gal80<sup>ts</sup>;OK107/+* | 0.5740 ± 0.03641 | 0.3500 ± 0.06351 | 0.2017 ± 0.08769 |
| *Gal80<sup>ts</sup>;OK107/RNAi-Syt1* | 0.6700 ± 0.04487 | 0.3733 ± 0.1032 | 0.4433 ± 0.1279 |
| *+ /RNAi-Syt1* | 0.6140 ± 0.03957 | 0.3983 ± 0.08118 | 0.6750 ± 0.06270 |

